# An alignment-last approach enables rapid transcriptomic biomarker discovery in large cohorts

**DOI:** 10.64898/2026.08.31.748301

**Authors:** Erkan Narmanli, Alexandre Lanau, Mara Neacsu, Mariia K. Koshkina, Pierre Fumeron, Philippe Martin, Nicolas Servant, Nicolas Perrin-Gilbert, Joshua J. Waterfall

**Affiliations:** Inserm U1330, Institut Curie Centre de Recherche, PSL University, Paris, France; Translational Research Department, Institut Curie Centre de Recherche, PSL University, Paris, France; Computational Oncology, PSL Research University, Mines Paris Tech, INSERM U1331, Paris, France; Institut des Systèmes Intelligents et de Robotique (ISIR), Sorbonne Université, CNRS, Paris, F-75005, France

## Abstract

Canonical transcriptomic analysis requires committing from the outset to a reference genome or transcriptome, which imposes a predefined feature set, usually annotated genes or isoforms. Alignment and annotation dilute the signal through feature-level aggregation, discard any sequence absent from the reference, and require reprocessing the entire dataset for each new question (mutations, fusions, transposable elements). Here, we introduce the *alignment-last* paradigm, in which the read becomes the unit of comparison across samples, and alignment is deferred to annotate only the relevant sequences. Querying the *merome*, a reference-free cohort k-mer index, with just a handful of reads (about 0.01% of a sample’s) reveals the cohort’s transcriptomic structure in bulk and single-cell data. At single-cell resolution, these reads outperform genes for cell classification and rediscover, without supervision, a transposable-element signature (VL30) of exhausted T cells. Finally, unsupervised read-level differential analysis recovers established lncRNA biomarkers; uncovers new prognostic transposable-element reads in adrenocortical carcinoma and sarcomas; and extracts signals even from reads that fail to align.

## Results

Canonical RNA-seq analysis follows a standardized structure (sequencing, quality control, alignment against a reference genome or transcriptome, quantification, differential analysis) in which alignment plays a structurally dominant role, both in computational cost and in the methodological choices it imposes downstream. This dominance has three limiting consequences. First, alignment is lossy: unalignable reads (exogenous viral sequences, somatic retrotransposon insertions^1,2^, atypical junctions) are filtered out, multi-mappers arbitrarily assigned or removed^3^, and any signal below quality thresholds is discarded. Second, alignment is annotation-dependent: each read is informative only against a reference, and the very unit of comparison between samples (typically the gene) is fixed by this annotation, biasing analysis toward the known transcriptome and marginalizing non-coding elements^4^, atypical ORFs^5^ and structural variants^6^. Third, alignment cost is not amortized: no single run covers all future questions, and each new inquiry (point variants, gene fusions, viral detection, transposable-element quantification) demands its own alignment. At the scale of large cohorts (e.g. TCGA and GTEx totaling approximately 170 TB of FASTQ), a complete realignment exceeds a year of CPU time and is beyond a standard laboratory; even Recount3^7^, a consortium effort for uniform re-quantification, locks cohorts into a single alignment run constrained by its annotation choices.

The size of these cohorts is therefore both their strength and their limitation: their scientific value erodes once analysis stays confined to the annotations chosen at publication, and any deep reanalysis meets a wall of computational cost. Several families of alignment-free methods have each addressed part of this problem over the past decade. The first family targeted the cost of expression quantification: Kallisto^8^ and Salmon^9^ introduced pseudoalignment, rapidly estimating transcript abundance by assigning each read to compatible transcripts without base-by-base alignment; now standard for quantifying individual samples, they nonetheless remain tied to a fixed transcriptomic reference and offer no native mechanism for cohort-wide analysis or annotation-independent discovery.

A second family of methods indexes k-mers at whole-cohort scale. Sequence Bloom Tree^10^ returns a presence/absence status from per-sample Bloom filters, while quantitative variants add abundance estimation: REINDEER’s monotigs^11^, MetaGraph’s counting de Bruijn graphs^12,13^ and Needle’s interleaved Bloom filters^14^. These make cohort-scale indexing accessible but leave the choice of query sequences to the user, embed no framework for differential analysis or classification, and build no common, data-derived reference for comparing samples.

A third family builds such a reference directly from the cohort’s k-mers: DE-kupl^15^ and kmdiff^16^ test each k-mer for differential abundance across conditions from raw reads, with DE-kupl additionally assembling significant k-mers into genome-aligned contigs; SPLASH2^17^ instead flags sites whose sequence composition differs between samples. All analyse without a reference and align only a posteriori to annotate significant signals, an approach conceptually close to the *alignment-last* paradigm we formalize here; yet none builds a persistent cohortwide index: DE-kupl and kmdiff scale to only a few dozen samples, and SPLASH infers without a reusable index. No current method therefore combines (i) cohort-scale indexing, (ii) reference-free differential analysis and (iii) a sub-gene-level, data-derived reference for inter-sample comparison.

Here we introduce *alignment-last*, a method that performs quantification and differential analysis directly on raw reads and defers alignment to a final annotation step, reversing the standard pipeline order. Its central idea is to replace conventional expression features (gene, transcript, exon) with the read itself as the shared unit of analysis across samples; a panel of reads sampled from the cohort’s FASTQ files is queried against all samples to fuel classification or differential analysis. The read is thus both the unit of observation and the axis of inter-sample aggregation, preserving sub-gene resolution while enabling direct cohort-wide comparison without materializing the exhaustive k-mer matrix. To make this operational we build the *merome*, the integrative collection of a cohort’s read repertoires indexed by an *interleaved Bloom filter* ^18,19^ of minimizers^20^; our implementation uses Needle^14^ for its sensitivity to low-abundance signals, though any cohort-scale backend with an equivalent quantitative property would serve.

We deploy this paradigm on a pan-cancer and a pan-tissue cohort, TCGA and GTEx (more than 28,000 samples), and extend it to single-cell RNA-seq (SMART-Seq and 10x). Beyond recovering known biomarkers and matching gene-level quantification, read-based analysis turns these cohorts into a discovery platform. When the abundance of randomly sampled reads is used to train a classifier, 1,000 reads suffice to distinguish 33 tumor types at 91.8% accuracy. Classifiers trained on reads even outperform the ones trained on genes when sampling less than 100 features In single cells, the merome outperforms gene-level quantification (67% versus 60% accuracy, F1 score 0.64 versus 0.57), and from random reads alone it recovers a VL30 transposable-element signature of exhausted T cells established by a dedicated multi-omic study^21^. More strikingly, read-level differential analysis between tumors and normal tissues rediscovers known oncogenic lncRNAs without prior annotation and at read resolution, each in its expected tumor type and replicated in the independent CPTAC cohort. Without supervision, the same analysis uncovers transposable-element reads whose expression stratifies patient survival. An LTR10D read marks better outcome in adrenocortical carcinoma (log-rank *P* = 0.0009), and an L1ME2z read marks poorer outcome in sarcoma (log-rank *P* = 0.0058), where it is detected in 69% of sarcomas versus 11% of healthy tissue. Alignment-last thus exploits a fraction of the transcriptional signal that gene aggregation dilutes and reference alignment discards.

### Reads as a fast and reference-free cohort comparison unit

The *alignment-last* differential analysis paradigm relies on a cohort index we call the *merome* (from *µɛ́ρoς méros*, “fragment”, and the suffix -*ωµα* -*ōma*, “totality”), in which the *k* -mer, rather than the read, is the unit of indexing. Querying a merome with a sequence returns one abundance estimate per sample, normalized by the sample’s sequencing depth so that estimates are comparable across the cohort. Because a query is matched through the kmers it contains rather than as a whole read, this does not require an exact, end-to-end correspondence with a cohort sequence.

The pipeline (Fig. 1a) keeps the standard components of transcriptomic analysis (alignment, annotation, quantification, and differential analysis) but rearranges their order. Samples are preprocessed without alignment (quality control and trimming only), then indexed into a merome at a one-time cost amortized across all later analyses. The expression matrix is built by querying the merome: reads randomly drawn from samples of interest each return an abundance vector across the cohort. This reads-by-samples matrix feeds a classifier or a differential analysis across conditions, which returns differentially expressed *reads* rather than *genes*. This resolution preserves sub-gene signals (point variants, atypical splicing, transposable-element exonization, gene fusions) otherwise lost to gene aggregation. Only these reads of interest are finally aligned (e.g. by STAR^22^) and annotated (e.g. by HOMER^23^: gene, genomic region, transposable-element class, distance to the TSS, etc.), so the read stays the unit of analysis until characterization, without the transcript or splice-site reconstruction that conventional pipelines perform by local assembly after alignment. Performed downstream, alignment and annotation no longer generate the biological signal but interpret it. It also frees annotation choices (reference genome, variant catalog, viral set, splicing models) from being prerequisites: rather than applied to the whole cohort upstream, they follow the reads the analysis actually selects.

**Fig. 1.**
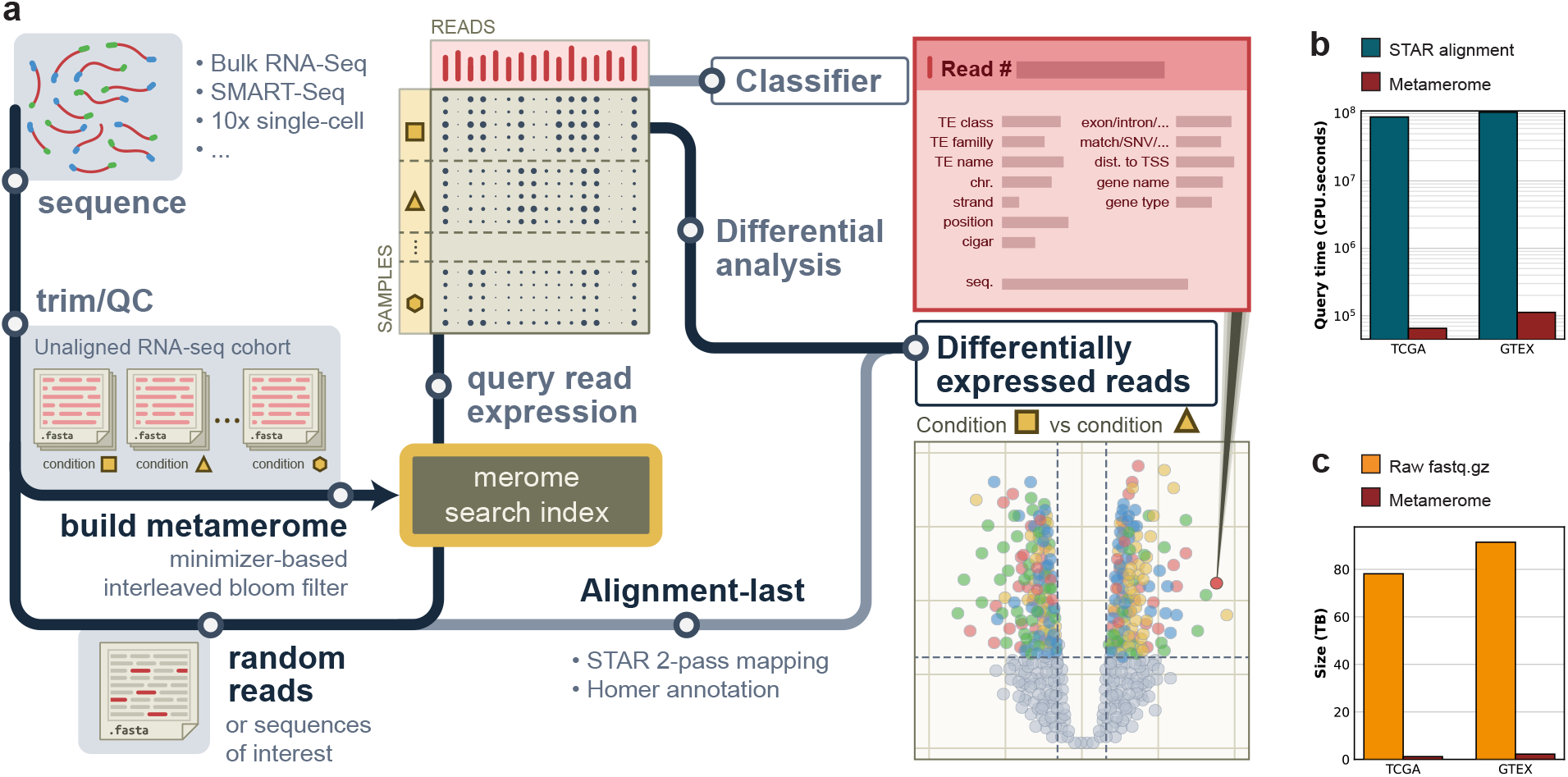
Transcriptomic meromes introduce a new paradigm for alignment-last differential analysis. Overview of the *alignment-last* workflow: reads or sequences of interest are queried against the merome (the integrative collection of the *k* -mers of a cohort) to build an expression matrix, which feeds classification or differential analysis; only the selected reads are then aligned and annotated. **b**, Computation time to query the reference transcript sequences of the 63,187 Gencode v45 genes against the TCGA and GTEx meromes (red) versus conventional genome alignment (blue); CPU-seconds, log scale. **c**, Size of the merome indices (red) versus the raw sequencing data (fastq.gz, orange), for TCGA and GTEx.

This reordering rests on a single cohort-wide index: built once per cohort, these meromes serve as reference resources throughout this study (TCGA^24^, GTEx^25^; Fig. S1). A merome accepts any query sequence, such as a single read, a variant, or a full-length transcript. Querying these indices with the full reference transcript sequences of the 63,187 Gencode v45 genes is about three orders of magnitude faster than the equivalent STAR alignment (Fig. 1b), and the indices are about two orders of magnitude more compact than the raw sequencing data (Fig. 1c). Because the merome is built once and thereafter only queried, its construction cost (*∼*2 min.CPU/GB) is amortized from the first analysis onward and any subsequent re-query is effectively free (*∼*0.21 min.CPU/Mbp/sample), making the exploration of cohorts that span hundreds of terabytes tractable on a standard workstation within hours. Yet computational performance and compactness do not guarantee accuracy: minimizer sampling and the merome’s probabilistic nature must be validated against biological observables.

### Meromes recover tumor biomarkers and cohort structure

We first tested merome estimates on biomarkers absent from reference genomes, which conventional alignment struggles to characterize, then confirmed that they reproduce STAR gene quantification and preserve each cohort’s transcriptomic structure.

Human papillomaviruses (HPV) set our first test case: their E6 and E7 oncogenes drive HPV-induced cancers^29^, yet are absent from the human reference genome, and therefore fall outside the scope of conventional pipelines. We queried these genes for the three most prevalent HPV strains^26^ against the TCGA meromes. HPV16 E7 (Fig. 2a) is most often positive in cervical (CESC) and head and neck (HNSC) cancers, two established HPV16 tropisms^30,31^. Merome-positive but reference-negative samples concentrate in the lower tail of the normalized expression distribution, consistent with detection below the reference calling threshold; across all 33 TCGA projects (Fig. 2c, HPV section), the F1 score against this reference stays high for all four strains.

**Fig. 2.**
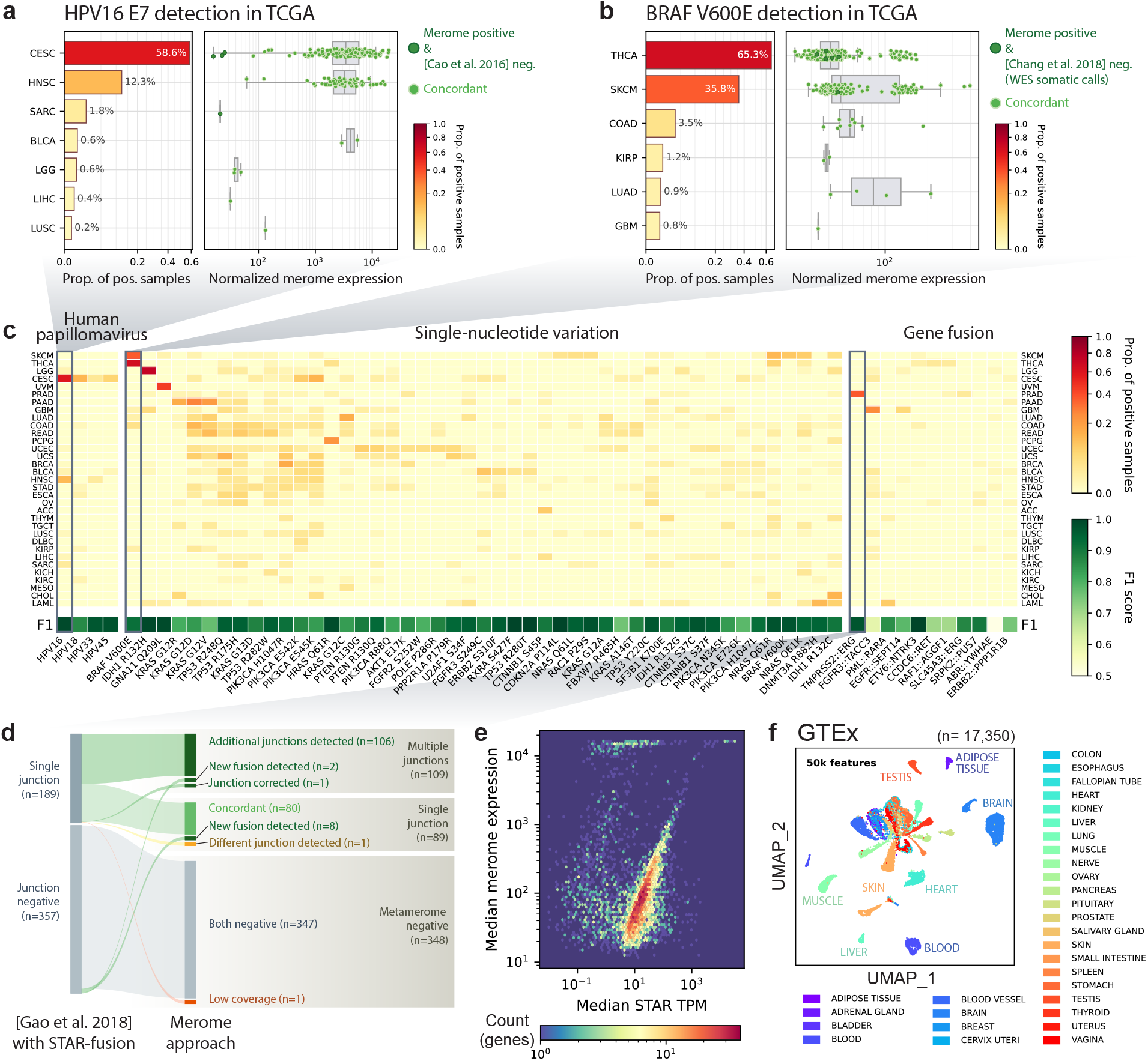
Meromes faithfully reproduce known biology: known-biomarker detection, reference gene quantification, and cohort structure preservation. **a**, Merome detection of HPV16 E7 across tumors (TCGA). Left: proportion of positive samples per project (square-root scale). Right: normalized expression among positive samples. **b**, Merome detection of BRAF V600E across tumors. Left: proportion of positive samples carrying the mutation (square-root scale). Right: normalized expression among positive samples. **c**, Most common reference biomarkers across the 33 TCGA projects (proportion of merome-positive samples, square-root scale): human papillomavirus ^26^ (E6/E7 genes), SNV hotspots ^27^ and gene fusions ^28^. Each category includes only biomarkers with at least 10 positive samples: across all projects for oncoviruses and fusions, or within a single project for SNV hotspots. Bottom row reports the F1 score between the reference call-set and merome detection. Per-biomarker proportions and F1 scores are in **Supplementary Data 1**. **d**, Merome versus STAR-Fusion detection of the TMPRSS2::ERG fusion in prostate cancer. The 26-bp junction-symmetric queries recover known junctions (*n* = 80 concordant), new ones (*n* = 10) and additional junctions in already-positive cases (*n* = 106). **e**, Median gene expression (Gencode v45), quantified by STAR (TPM) versus the merome (querying each gene’s full RNA sequence); hexbin, log scale. Spearman *ρ* = 0.57 for typical expression regime (STAR TPM *≥* 5; 78% of co-detected genes); Spearman *ρ* = *−*0.16 for very low expression regime (STAR TPM *<* 5; 22%). Per-gene medians are in **Supplementary Data 2**. **f**, UMAP of the 17,350 GTEx samples from a random panel of 50,000 reads; samples cluster by tissue.

To test the merome’s nucleotide resolution we queried pan-cancer SNV hotspots ^27^ as paired reference-and-variant 25-bp sequences (Methods). Across all 33 TCGA projects (Fig. 2c, SNV section), the F1 score against the cBioPortal call-sets^32,33^ exceeds 0.8 for nearly all hotspots, and the expected tissue specificity is preserved: e.g. BRAF V600E concentrated in SKCM^34^ and THCA^35^, KRAS mutations^36^ in PAAD, LUAD, COAD and READ. BRAF V600E is the activating mutation found in roughly two-thirds of melanomas^34^ and is targeted by selective inhibitors^37^; our meromes detect it in SKCM and THCA (where it marks papillary variants) but not significantly elsewhere (Fig. 2b, Fig. S2a,b). Because this reference call-set is exome-based^32,33^ whereas the merome measures transcriptomic expression, some discordance is expected; as for HPV, the reference-negative variants concentrate in the lower tail of the expression distribution, consistent with greater sensitivity to low-abundance variants.

A gene fusion creates a chimeric junction sequence, absent from the reference genome, whose *de novo* detection by alignment requires dedicated pipelines (STAR-Fusion^38^, Arriba^39^) that reconstruct junctions from chimeric reads. Merome does not serve this exploratory role but allows targeted interrogation of suspected fusions from their junction sequences (Methods), complementary to discovery tools. Querying the reported junctions^28^ for the canonical TMPRSS2::ERG fusion in TCGA-PRAD^40^ retrieves 188 of the 189 reference-positive cases, recovers the fusion in 10 reference-negative samples, and detects additional junctions over those identified by STAR-Fusion in 106 cases (Fig. 2d, Fig. S2c). Across the fusion heatmap (Fig. 2c, **Supplementary Data 1**) the merome recovers all documented fusions with high F1 for most (*F* 1 > 0.69 for 8 of 10 fusions), completing binary validation across the three biomarker classes.

Beyond non-reference sequences, we verified that merome estimates reproduce gene-level STAR expression (Fig. 2e). Both approaches agree at typical expression levels (*ρ* = 0.57) but diverge in the very-low-expression regime, near the detection limit, where both quantifications are dominated by counting noise (Methods). Residual off-diagonal deviations are mostly genes overestimated by the merome, reflecting minimizer multi-mapping among paralogs, pseudogenes and repeats, an expected consequence of *k* -mer indexing, which does not resolve the ambiguity that alignment would (mean normalized sequence entropy of 10-mers: 0.41 off-diagonal vs 0.50 rest; t-test p-value: 1.5×10-55). This gene-level accuracy validates standard differential comparisons on merome expression; it remains to confirm that meromes also capture each cohort’s overall transcriptomic structure.

So far, the merome has been queried with known sequences of interest (genes, variants, junctions); to agnostically probe overall transcriptomic structure we now query reads. In all following analyses, reads serve as the unit of comparison. First, a random panel of 50,000 reads from the cohort is queried against every sample (Methods). The resulting reads-by-samples matrix, embedded by UMAP (Fig. 2f, S3), clusters by tissue for GTEx and by tumor type for TCGA, with the expected groupings (e.g. gastrointestinal cancers grouped together). Thus the transcriptomic structure of both healthy tissues and tumors emerges from a random *∼*0.00001% of the cohort’s reads, and the separation sharpens steadily with the number sampled (Fig. S3). That so few reads already reveal this structure without supervision raises the next question: how many are needed for reliable supervised classification?

### A handful of reads allow sample classification

Using reads as features in a supervised multi-class classifier (Methods), we ask how many randomly selected reads are needed to identify a tissue or tumor type. We build classifiers with different numbers of randomly selected reads (from 1 to 50,000), at each level performing 100 independent read samplings. We report summary statistics over the independent read draws as indicated for robustness. A parallel analysis on gene quantification (STAR GeneCounts TPM) directly compares reads and genes as units of information.

At 1,000 reads, the classifiers reach 91.8% accuracy on TCGA (Fig. 3d) and 96.9% on GTEx (Fig. S4d). Residual misclassifications concentrate among biologically related tissues (e.g. colon, esophagus, bladder, all epithelial) rather than unrelated ones. The TCGA glioblastoma set (TCGA-GBM, 31 samples, Fig. 3a) illustrates a harder-to-classify tumor type (a healthy-tissue case is in Fig. S4a–c). Per-sample accuracy rises monotonically with features: 25 of 31 samples are robustly classified at 50 reads, 30 at 1,000. At 50 reads (Fig. 3b,c), misclassified samples mostly alternate with a low-grade glioma (LGG) call before converging to the right label; one atypical sample is classified as sarcoma even at 50,000 reads. Using reads as features directly raises the question of their informational efficiency versus gene quantifications.

**Fig. 3.**
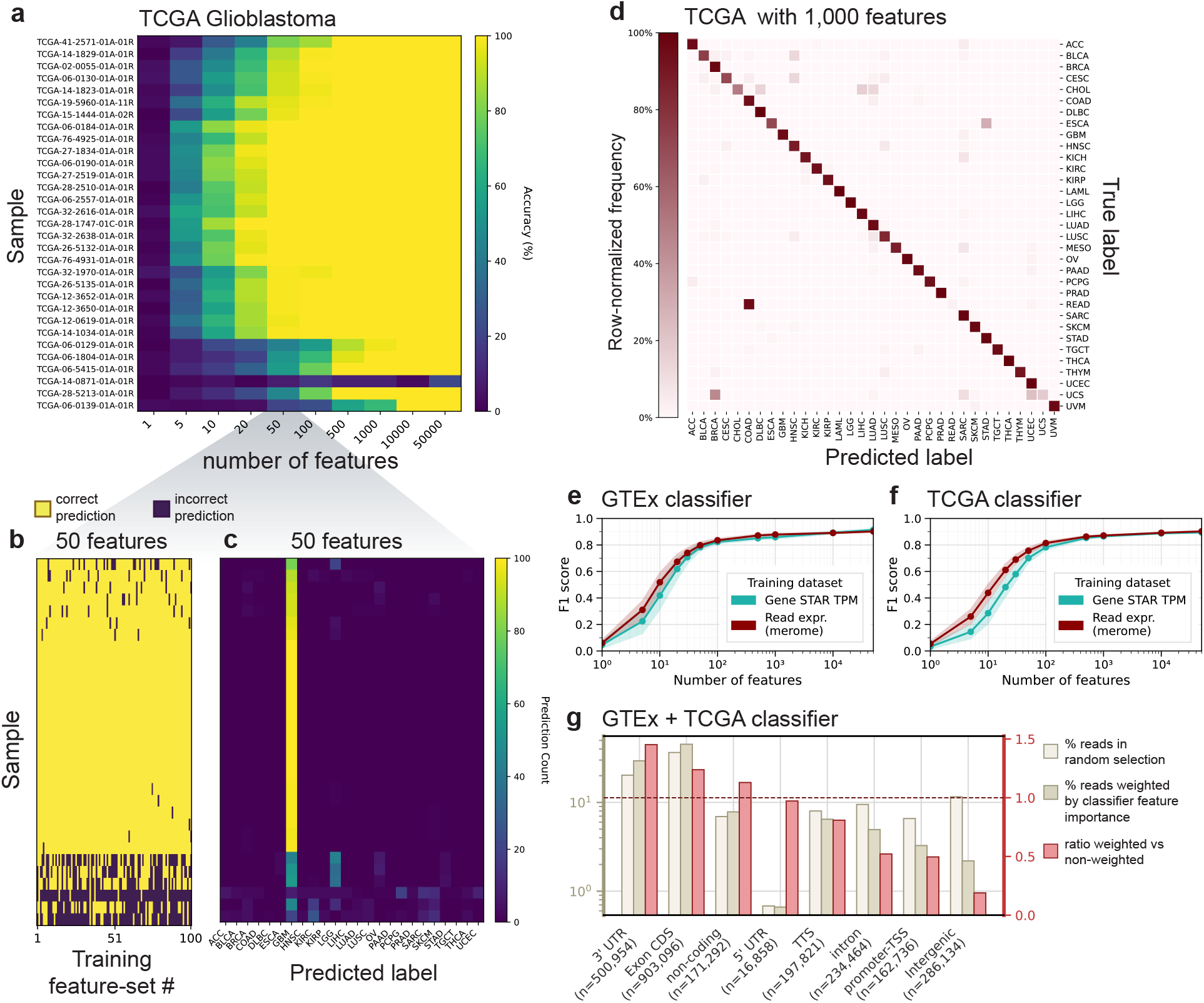
A handful of random reads classifies transcriptomic samples with accuracy comparable to gene-expression methods. **a**, Multiclass tumor classification (33 tumors) of true-labeled glioblastoma (GBM) samples (TCGA-GBM; 31 test samples) versus number of features. Each row is a sample, each column a feature-set size (random reads; 1 to 50,000); color, proportion of correct predictions over 100 independent draws. Samples ordered by hierarchical clustering. **b**, At 50 features, prediction per sample (rows) and feature-set (columns; 100) on the GBM test set. **c**, At 50 features, distribution of predicted labels (columns) per GBM test sample (rows); color, number of draws (of 100) assigning each label. **d**, Confusion matrix for the TCGA classifier (reads, 1,000 features; mean of 100 runs; overall accuracy 91.8%). **e**, **f**, F1 score versus number of features for the GTEx (**e**, 30 tissue types) and TCGA (**f** ) classifiers, using reads (merome; red) or genes (STAR TPM; blue); mean *±* s.d. over 100 runs. **g**, Informational enrichment of genomic regions for the joint GTEx+TCGA classifier (63 classes; 50,000 features; 50 runs). Per HOMER region (3’UTR, CDS exon, non-coding, 5’UTR, TTS, intron, promoter-TSS, intergenic): proportion of reads in the random sampling (light beige), the Random-Forest-importance-weighted proportion (dark beige), and their ratio (red; right axis). Dotted line, ratio of 1 (no enrichment).

Calculating the F1 score as a function of the number of features for each modality (Fig. 3e,f) directly weighs the information in reads against genes. Beyond 1,000 features the two converge on GTEx and TCGA: reads and genes then carry the same discriminatory power. Yet a gene aggregates hundreds to tens of thousands of aligned reads; despite this asymmetry, below 100 features reads outperform genes, a few dozen random reads rivaling gene expression for discriminating tissue and tumor types. This demonstrates the advantage of sub-gene resolution: a read may land on a tissue-informative splice junction ^41^ or a tumor-informative TE fragment^42^, signals that gene aggregation dilutes into a count average. In a low-feature-count setting, then, *resolution* (sub-gene versus gene) outweighs *volume*.

To characterize the reads carrying the discriminating signal, we weighted each read’s genomic annotation by its importance in the joint GTEx+TCGA classifier and compared the importance-weighted composition with the raw read composition (Fig. 3g and Methods). Exonic regions (specifically 3’UTRs and CDS exons) show the highest ratios, so the discriminating signal is dominated by the coding transcriptome. Annotated non-coding RNAs also contribute importantly to classifier performance, consistent with their high tissue specificity^43^. Introns, intergenic regions and promoters show low overall specificity, though individual loci can be highly informative in context^42,44^.

Multiclass classification thus quantitatively validates the information carried by meromes: 1,000 reads suffice to achieve high performance, and individual reads outperform gene quantification even on a small feature budget. It reflects the richness of the read’s sub-gene signal (coding and non-coding) that gene aggregation dilutes and reference alignment filters out. We pursue this sub-genomic richness next in single-cell RNA-seq.

### Single-cell meromes outperform alignment-first approach and rediscover TE biomarkers

Single-cell RNA-seq challenges the merome approach for several reasons, including fewer reads per sample, end-capture biases (3’ or 5’ for 10x^45^), and cell subpopulations more transcriptomically alike than heterogeneous tissues. We test it on two complementary technologies: SMART-Seq (full-length) and 10x (3’ or 5’ biased), showing that single-cell meromes classify cell types and, from random reads alone, rediscover without supervision a transposable-element signature of exhausted T cells.

SMART-Seq generates full-transcript reads, a bulk-like profile well-suited to merome analysis. We use a SMART-Seq dataset of mouse thymic development^46^ where cells were FACS-sorted by surface phenotype, so the labels are protein markers independent of the transcriptomic data. In the thymus, CD4^+^CD8^+^ double-positive (DP) progenitors differentiate into CD4 or CD8 single-positive (SP) cells via selection intermediates (CD69*^−^* DP, CD69^+^ DP, TCR^hi^ DP, CD4^+^CD8^low^) forming a transcriptomic continuum. UMAPs from 10,000 features (Fig. 4a,b) cluster most states (except TCR^hi^ DP) but do not fully separate the six subpopulations: with both methods, the selection-intermediate continuum remains hard to resolve, as expected. In contrast, the merome separates CD4 SP from CD8 SP better than STAR TPM. Random Forest classification quantitatively confirms this advantage, the merome outperforming STAR TPM in both accuracy (67% versus 60%) and F1 (0.64 versus 0.57; Fig. 4d-e). The confusion matrices show a clearer diagonal for the merome, with fewer misclassifications between intermediate subpopulations (Fig. 4c, S5a,b). The merome advantage seen in bulk thus holds in SMART-Seq single cells.

**Fig. 4.**
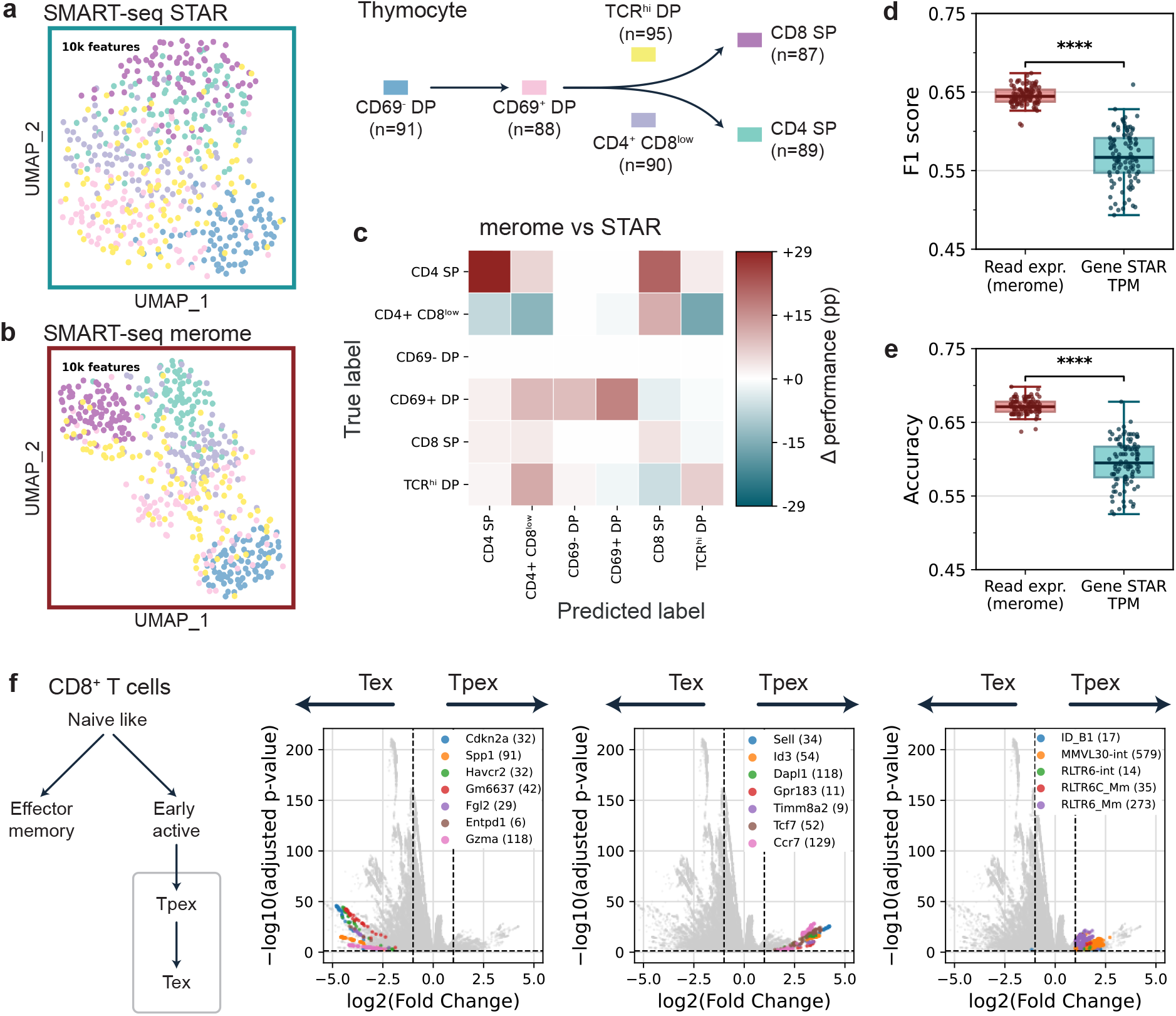
Single-cell meromes outperform conventional gene quantification for cell classification and reveal transposable-element signatures in exhausted T cells. **a-b**, UMAP of mouse thymocytes (SMART-Seq, ^46^) on 10,000 features, from STAR TPM (**a**) and merome (**b**). Cells were FACS-sorted by surface phenotype (anti-CD4, CD8, CD69, TCR*β*), giving labels independent of the transcriptome. Six thymic subpopulations: CD69*^−^* DP progenitors (CD4^+^CD8^+^CD69*^−^*, *n* = 91); selection-intermediate CD69^+^ DP (CD4^+^CD8^+^CD69^+^, *n* = 88), CD4^+^CD8^low^ (*n* = 90), TCR^hi^ DP (CD4^+^CD8^+^TCR*β*^hi^, *n* = 95); and differentiated CD4 SP (CD4^+^CD8*^−^*, *n* = 89), CD8 SP (CD4*^−^*CD8^+^TCR*β*^+^, *n* = 87). **c**, Term-by-term difference between the merome and STAR-TPM Random Forest confusion matrices on SMART-Seq thymocytes (10,000 features, 100 runs); color codes the merome advantage in percentage points (Methods). Overall accuracy 67.0% (merome) versus 60.2% (STAR-TPM); the two matrices appear separately in Fig. S5a,b. **d-e**, Box plots of F1 (**d**) and accuracy (**e**) (10,000 features, 100 runs) for reads (merome: median *F* 1 = 0.6445, accuracy=67.2%) versus genes (STAR TPM: median *F* 1 = 0.5669, accuracy=59.5%) on the SMART-Seq thymus dataset (Student’s t-test: F1 *p* = 1.35*e −* 55; accuracy *p* = 4.61*e −* 59). **f**, Left: CD8^+^ T-cell differentiation scheme. Right: three volcano plots of reads differentially expressed between Tpex (Slamf6^+^) and Tex (Tim3^+^). Genes and TE subfamilies are ranked by the mean log2FC of their differentially expressed reads, keeping only those with *≥* 5 such reads. The first two plots highlight the 7 top genes in Tex and Tpex; the third, the 5 top TE subfamilies in Tpex (MMVL30-int, RLTR6_Mm, RLTR6C_Mm, RLTR6-int, ID_B1), matching previously identified TE signatures ^21^.

The second dataset comes from 10x Genomics, the most widely used scRNA-seq technology, and addresses CD8^+^ T-cell exhaustion^21^. Exhausted T cells are heterogeneous^47^, comprising early progenitors (Tpex, Slamf6^+^) and terminally exhausted cells (Tex, Tim3^+^); unlike Tex, Tpex retain proliferative potential and respond to anti-PD-1^48^. Here, 10x captures only the 5’ end of transcripts^45^, reducing accessible sequence diversity, a bias inherently unfavorable to k-mer indexing. On this dataset the goal is not classification but read-level differential analysis: adopting the published Tpex/Tex labels^21^, we set out to reproduce that study’s central result, a Tpex-specific enrichment of VL30 transposable elements it established with a dedicated multi-omic pipeline (bulk RNA-seq, RT-qPCR and ATAC-seq). We instead query only random reads against a 10x merome, without any prior transposable-element annotation.

Read-level differential analysis between Tpex and Tex produces volcano plots (Fig. 4f and Methods) with a distinctive shape: differentially expressed reads form “trails” of aligned points, each trail a set of reads from the same locus (a signal usually aggregated into one gene, here resolved read by read). Annotating these reads (alignment-last) identifies the expected canonical markers (Fig. 4f left, center; Fig. S5c,d for the top 20). In Tex cells, top genes include *Havcr2* (TIM3), an inhibitory receptor of the most terminally exhausted cells^49^, and *Cdkn2a*, a cell-cycle inhibitor. In Tpex, *Tcf7* (TCF1), a transcription factor of exhausted progenitors^48,50^, and *Sell* (CD62L), a memory-T-cell homing receptor. Among the top upregulated TE subfamilies in Tpex (Fig. 4f, right; Fig. S5e), two stand out, MMVL30-int and RLTR6_Mm, both VL30 LTR retrotransposons. Without supervision, and from random reads alone, this recovers the same Tpex-specific VL30 enrichment, and gives an independent validation of that study’s central result.

### Differentially expressed reads uncover unconventional cancer biomarkers

Guided by the enrichment of non-coding reads in classification (enrichment ratio *>* 1; Fig. 3g), we applied read-level differential analysis to the non-coding transcriptome in the TCGA and GTEx cohorts. As before, reads sampled across both cohorts are queried against the meromes, each tumor project is contrasted with all other samples (tumours and healthy tissues) without prior annotation, and significant reads are aligned and annotated a posteriori. Retaining the reads overexpressed in each tumor and annotated as non-coding yields 391 lncRNA reads (Methods). As expected from their selection, these separate the 11,137 TCGA tumors by type (Fig. 5a); notably, they also organize the 17,350 GTEx samples by tissue (Fig. S6a).

**Fig. 5.**
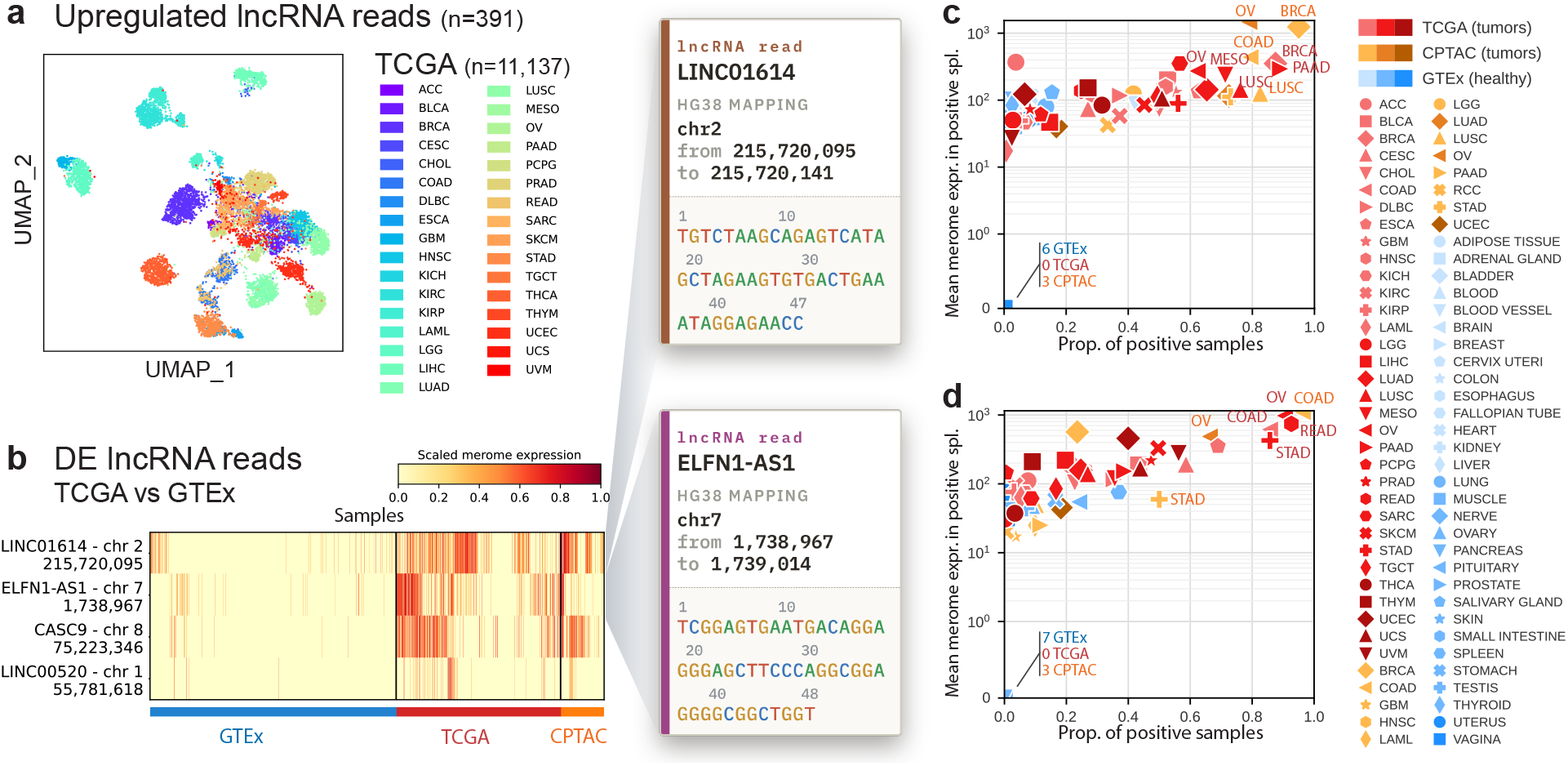
Upregulated lncRNA reads in cancers distinguish tumor cohorts; rediscovery of established oncogenic lncRNA biomarkers. **a**, UMAP of TCGA tumor samples from the 391 tumor-upregulated lncRNA reads (**Supplementary Data 3**) identified by TCGA-versus-GTEx read-level differential analysis; each point is a sample, colored by tumor project. **b**, The four lncRNA reads exceeding log2FC *≥* 4 between TCGA and GTEx: LINC01614 (chr2), ELFN1-AS1 (chr7), CASC9 (chr8) and LINC00520 (chr14). Heatmap of their merome expression, min–max-scaled 0–1 per read; each column is a sample, grouped by cohort: GTEx (healthy), TCGA (tumors, discovery) and CPTAC (tumors, validation). **c**, LINC01614 read: proportion of positive samples (x-axis) versus mean merome expression among expressing samples (y-axis, symlog); each point is a project (TCGA and CPTAC tumors; GTEx healthy tissues). LINC01614 read was detected in fewer than 5% of samples in 18 of 30 GTEx tissues. **d**, ELFN1-AS1 read: as in **c**; detected in fewer than 5% of samples in 23 of 30 GTEx tissues.

Ranking these reads by their TCGA-versus-GTEx expression difference, four long non-coding RNAs (lncRNAs) stand out (log2 fold change above 4, Fig. 5b): LINC01614, ELFN1-AS1, CASC9 and LINC00520. These four reads remain nearly unexpressed across most healthy tissues (Fig. 5c,d; Fig. S6c,d), and their tumor-specific overexpression is fully replicated in the independent CPTAC cohort^51^, ruling out a discovery-cohort artifact. Each read is overexpressed precisely where its lncRNA is already described as oncogenic: LINC01614, a poor-prognosis biomarker in breast cancer ^52^; ELFN1-AS1, oncogenic in colorectal and ovarian cancers^53,54^; CASC9, upregulated in head-and-neck and lung squamous carcinomas^55,56^ and the most overexpressed lncRNA in its esophageal squamous-carcinoma screen^57^; and finally LINC00520, oncogenic in head-and-neck squamous carcinoma^58^ and melanoma^59^. The approach thus rediscovers four established biomarkers at single-read resolution, each in its expected tumor type, without prior annotation. Several reads also flag expression in tumors not yet characterized for these lncRNAs: for example, LINC01614 is detected in 71% of mesotheliomas and 62% of ovarian cancers, and CASC9 in 75% of testicular germ-cell tumors, consistent with a very recent study ^60^. The same read-level analysis next turns to transposable elements and, finally, to unaligned reads.

We then examined the overexpressed reads in each tumor that align to TEs (Fig. S7; **Supplementary Data 4 and 5**). Two stand out: the LTR10D element (LTR/ERV1) in adrenocortical carcinoma (ACC) and the L1ME2z element (LINE-1) in sarcoma (SARC). The LTR10D dup158 read is overexpressed in 38% of ACCs, above other tumors and healthy tissues (Fig. 6a,b), and its presence is associated with better overall survival (Fig. 6c and Methods). Positive samples concentrate in the favorable-prognosis ACC subtypes (COC1, low-steroid phenotype, C1B) and are rarer in aggressive ones, consistent across the three reference classifications^61^ (Fig. 6d, S8 and Methods). The full-length element reproduces these enrichments and the survival association, confirming the read faithfully reports it (Fig. S8 and Methods). Gene-level annotation also recovers the established ACC biomarker miR-483, an intronic microRNA of IGF2^62,63^.

**Fig. 6.**
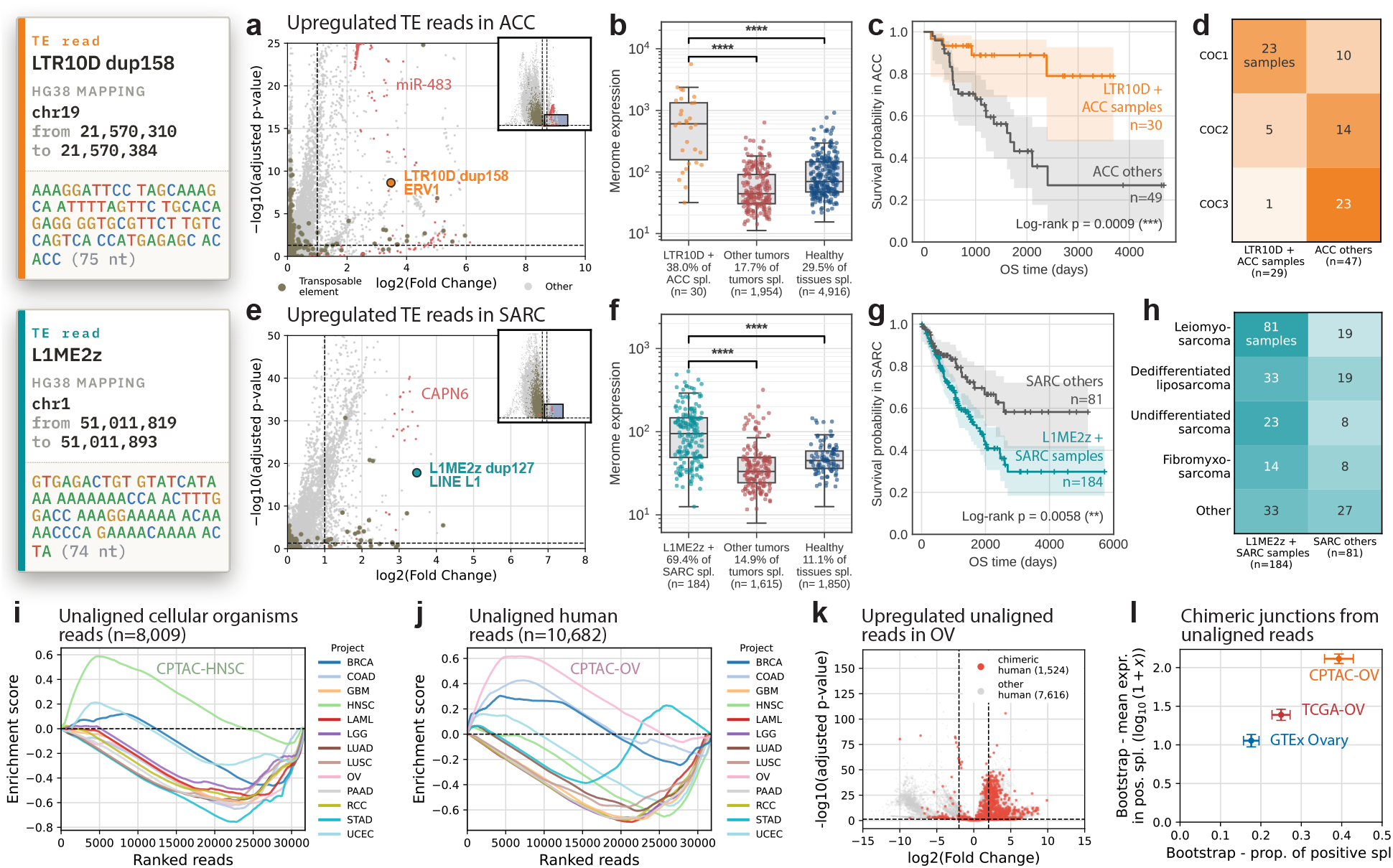
Differentially expressed reads allow unsupervised discovery of TE-derived and unaligned biomarkers. **a**, Upregulated reads in ACC (TCGA-ACC versus the rest: other TCGA tumors and GTEx healthy tissues); the LTR10D dup158 read (ERV1/LTR, orange) and reads aligned to the known miR-483^62,63^ (red; 650 reads) are highlighted. **b**, Merome expression of the LTR10D read in ACC versus other tumours and healthy tissues; higher in ACC (Mann–Whitney, *p* < 0.0001). **c**, Overall survival (Kaplan–Meier) in ACC, LTR10D^+^ samples versus others: read presence associates with better survival (log-rank *p* = 0.0009). **d**, LTR10D^+^ status versus the ACC “cluster-of-clusters” (COC) subtypes ^61^: enriched in COC1 (*z* = +4.96), depleted in COC3 (*z* = *−*4.14); *χ*^2^ *p* = 1.5*×*10*^−^*^6^, Cramér’s V 0.59. **e**, As in **a**, for sarcoma; the L1ME2z dup127 read (L1/LINE, turquoise) and reads aligned to the known CAPN6^64,65^ (red; 22 reads) are highlighted. **f**, As in **b**, for the L1ME2z read; higher in SARC (Mann–Whitney, *p* < 0.0001). **g**, As in **c**, for L1ME2z^+^ SARC samples; read presence associates with poorer survival (log-rank *p* = 0.0058). **h**, As in **d**, for L1ME2z^+^ status versus sarcoma histological subtype (subtypes with *≥* 15 samples, rarer ones pooled as “others”); present across all subtypes (leiomyosarcoma 81%, dedifferentiated liposarcoma 63%, undifferentiated sarcoma 74%, fibromyxosarcoma 64%, others 55%). **i**, Gene-set enrichment analysis (GSEA) of random CPTAC reads left unaligned and assigned to cellular organisms, ranked per project by the read-level differential statistic (one curve per project). **j**, As in **i** for unaligned reads assigned to the human genome. **k**, Volcano of unaligned human reads, CPTAC-OV versus other CPTAC tumours; reads spanning a STAR chimeric junction (red) versus other human reads (grey), with most chimeric reads upregulated in OV. **l**, For 26-bp queries reconstructed across chimeric breakpoints (1,332 junctions), the fraction of positive samples (x) and the mean expression among positive samples (y, log10) in CPTAC-OV, TCGA-OV (validation) and GTEx ovary (control), summarized by bootstrap over 10 resamples (point=mean; bars=*±* 2 s.d.).

The L1ME2z dup127 read, by contrast, associates with poorer survival (Fig. 6g), and is detected in 69% of sarcomas versus 15% of other tumors and 11% of healthy tissues (Fig. 6e,f). Unlike LTR10D, it spans all sarcoma histological subtypes (55–81% prevalence; Fig. 6h, S9). The read straddles two adjacent repeats, AluSz and L1ME2z; the sarcoma specificity and survival association trace to L1ME2z alone, AluSz being neither differentially expressed nor prognostic (Fig. S10 and Methods). Gene annotation likewise recovers CAPN6, a known oncogene in uterine leiomyosarcoma^64^ and liposarcoma^65^. Like the accompanying gene biomarkers, these two transposons were found without supervision. All reads analyzed so far align to the human genome; a final, unreferenced fraction of the tumor transcriptome is examined next.

We turned to the reads STAR leaves unaligned, which conventional pipelines discard, repeating the read-level differential analysis within CPTAC, each tumour project against the rest. We used CPTAC rather than TCGA, whose tumour taxonomic signal has proven largely artefactual, driven by contamination and analysis errors^66,67^. Ranking the unaligned reads of each project and testing them for enrichment by taxonomic origin^68^, we found reads assigned to cellular organisms enriched in head-and-neck (HNSC), colon (COAD) and breast (BRCA) carcinomas (Fig. 6i), cancers with an established tumour-associated or mucosal microbiota^69–72^. Reads assigned to the human genome yet left unaligned by STAR were instead most enriched in serous ovarian carcinoma (OV; Fig. 6j).

High-grade serous ovarian carcinoma is extensively structurally rearranged^73^, suggesting these unaligned human reads might mark structural variants. A dedicated STAR pass tuned for chimeric detection^39^ showed that, among the OV-upregulated unaligned human reads, the large majority of those spanning a chimeric junction were upregulated (Fig. 6k), as expected of rearrangement-derived reads. Reconstructing a 26-bp sequence across each candidate breakpoint and querying it back against the ovarian meromes, we found these junctions recurrently present in CPTAC-OV, reproduced in the independent TCGA-OV cohort, and less prevalent in healthy ovary (GTEx; Fig. 6l). Alignment-last thus recovers, from reads that conventional analysis discards, both a microbial signal and ovarian-specific structural-variant junctions.

## Discussion

In this study, we show that the read can serve as the unit of unsupervised biological discovery at the scale of the largest transcriptomic sequencing cohorts. Using randomly selected reads, we rediscover established oncogenic biomarkers and identify new non-coding elements associated with prognosis, not only in large tumor cohorts but also on single-cell data. The read thus becomes a new gateway to the transcriptome, opening up what alignment and annotation previously left out of reach.

Our aim here is to advance cohort k-mer indexes from validation toward discovery. Queried with chosen sequences, the *merome* recovers established biomarkers (Fig. 2), but reaches only those one already knows to look for; under the *alignment-last* paradigm, where reads are drawn at random, it instead surfaces biomarkers that were never specified in advance (Fig. 1). Classification and differential analysis are the two faces of this read-level view, the former isolating the reads a class shares as its consensus signature, the latter the reads that set it apart, both at the sub-gene resolution that gene aggregation would otherwise dilute.

To our knowledge, alignment-last is the first approach to unify a persistent cohort index, reference-free differential analysis, and a data-derived sub-gene unit of comparison, whereas earlier alignment-free methods each addressed only one piece of this problem^8,11,15^. Because the read is at once the index unit and the axis of comparison, a cohort indexed once can be re-queried at will for classification or unsupervised discovery, with alignment and annotation applied only a posteriori to the relevant reads.

Our approach has several limitations. First, the same minimizer may be shared by paralogs, pseudogenes, repeats, or regions of low complexity, which can lead to an overestimation of certain abundances without the disambiguation that an alignment would provide. Future work could anticipate this by precomputing, from a single pass over the reference genome, a library of minimizers that are not locus-specific, flagging upfront which counts call for disambiguation. Furthermore, the short read remains the unit of analysis, which prevents the analysis from capturing long-range relationships, such as the co-occurrence of events on the same transcript. Finally, the approach is not exhaustive; not all reads in a cohort are examined, and random sampling may overlook signals carried by reads that were never sampled. Detecting these would require upstream pre-filtering, for example via small meromes that select the most informative reads, an infrastructure that has yet to be built.

In short, the *alignment-last* approach turns the raw read into a unit of analysis in its own right, carrying the sub-gene signal that gene aggregation had previously masked. Its scope extends beyond the RNA-seq tumor cohorts studied here, as this paradigm could be applied to any collection of reads, regardless of their format. From the virome to metagenomics, fields lacking a complete reference genome would be the first to benefit; a version specifically tailored to RNA shotgun sequencing would then be particularly appropriate. Beyond differential analysis, the merome paradigm would accommodate other tools operating directly on the read, e.g. allowing candidate prioritization, for further immunogenicity assessment (peptide identification with immunopeptidomic, and in vitro/vivo immunogenicity assessments). To make these applications accessible, our Merome Explorer web platform will gradually expand its offering of indexed cohorts, building on the k-mer indexing efforts for public archives^13,74,75^. As these indexes and the methods that utilize them proliferate, sequencing repositories will cease to be static archives and become living resources that can be re-queried at will, freed from the constraints of alignment and annotation.

## Method

### Data and preprocessing

The TCGA data were downloaded using gdc-client v1.6.1 from the GDC manifests (as of January 2024). To limit RAM usage during merome creation, fastq.gz files larger than 13 GB were excluded, with the exception of six projects with large files (ESCA, GBM, LUAD, LUSC, OV, STAD), for which the threshold was raised to 23 GB, leaving 11,137 samples (98.79% of all samples, Fig. S1a). GTEx v8 data were downloaded via gen3-client from the AnVIL manifest (January 2024), segmented by tissue, with further subdivision by anatomical site for the five tissues exceeding 800 samples (adipose tissue, blood vessel, brain, esophagus, skin), resulting in 54 subprojects for 30 main tissues and 17,350 retained samples (Fig. S1b). The BAM files were converted to fastq using samtools sort followed by samtools fastq (samtools v1.9^76,77^). Read quality was assessed using FastQC v0.12.1^78^ and MultiQC v1.17^79^; trimming with trim_galore v0.6.9^80^ (built on cutadapt v4.2^81^) was applied. CPTAC RNA-seq data were downloaded from the GDC manifest (as of May 2026) with gdc-client v1.6.1 and split into 13 tumor projects, with all kidney pathologies grouped into a single RCC project. The CPTAC data were preprocessed exactly as the TCGA data.

The SMART-Seq dataset of mouse thymocytes^46^ and the 10x dataset of CD8^+^ TILs from B16-OVA tumors^82^ were used; the CD8^+^ TILs were annotated following the exhaustion study^21^. On the 10x data, we perfomd cell-quality filtering and Tex/Tpex annotation (Slamf6, Tim3 markers) with scanpy v1.11.0^83^ on alignments and count matrices generated by Cell Ranger v10^45^.

### Building and querying meromes

We indexed each cohort with Needle v1.0.1^14^ (needle ibf -w 25 -k 20 –cutoff 2 -l 10 - n 5 -f 0.05), which couples an interleaved Bloom filter (IBF)^18,19^ with minimizer-based sequence reduction^20^: for each sample, the IBF stores the presence of every minimizer over sliding windows. For bulk data, an index was built per project (33 TCGA, 54 GTEx) from concatenated paired-end fastq files. Only the CPTAC ^51^ meromes (13 tumor projects), added for the differential-expression analysis, were built later with the updated parameterization needle ibf -w 24 -k 18 –cutoff 2 -l 20 -n 5 -f 0.05. For single-cell data, an index was built per dataset, with each cell treated as an independent Needle sample. The read count per sample/cell was stored in metadata for normalization by sequencing depth.

Random read panels used as queries were drawn without replacement from the FASTQ files of the relevant samples, at the panel size and replicate draw specified for each analysis. Query sequences were grouped into batches of fewer than 100 million nucleotides and searched with needle estimate (Needle v1.0.3), which returns for each sample the median of the minimizer counts of the query. Because this median does not depend on the number of minimizers extracted, it is independent of query length, so length normalization is inherently applied. The resulting expression tables were converted into sparse matrices (scipy.sparse.csr_matrix; SciPy v1.15.1^84^) and encapsulated into AnnData objects (anndata v0.11.3^85^). Estimates were then divided by the total read count of the sample and multiplied by 10^9^ to obtain depth-normalized expression levels, applying the TPM normalization principle^86^ to make values comparable across samples.

### Quantitative assessment and resource benchmarking

The TCGA and GTEx fastq files were aligned against GRCh38.p14 (Gencode v45 ALL) using STAR v2.7.11b^22^ in –quantMode GeneCounts mode according to the ENCODE long-RNA-seq pipeline parameters^87^, and the raw counts were converted to TPM. In parallel, the 63,187 Gencode v45 gene sequences^88^ were queried against the TCGA and GTEx meromes, with estimates aggregated per gene.

Both pipelines were run single-threaded and timed by their workflow benchmark. For the merome, the per-index query CPU-seconds were measured directly and summed across the merome indices of each cohort (33 TCGA, 54 GTEx). For the STAR pipeline, the alignment of 15 fastq.gz files was extrapolated to the full cohorts via linear regression of CPU-seconds against the total fastq.gz size (*R*^2^ = 0.55, slope = 1.08 CPU*·*s/MB). Querying the 63,187 genes took *∼*18 and *∼*31 CPU-hours against the TCGA and GTEx meromes, respectively, versus *∼*1,027 and *∼*1,202 CPU-days for the corresponding STAR alignments (Fig. 1b); the indices compressed the cohorts from 78.1 to 1.2 TB (TCGA) and from 91.4 to 2.2 TB (GTEx), against *∼*170 TB of raw FASTQ in total (Fig. 1c). Two properties underlie this speed-up: the merome is pre-computed, so a query reads only the compact index rather than the cohort’s raw FASTQ files; and its interleaved Bloom filter tests each minimizer against the entire cohort in a single vectorized operation^14^, so that query cost scales with the length of the queried sequence rather than with the number of samples.

For each gene, the median over all TCGA and GTEx samples was computed for the depth-normalized merome estimate and for the STAR TPM, and the two estimates were compared on a hexbin grid (gridsize = 75, at least one gene per bin, log-scaled colour) over logarithmic axes, which display the genes with a positive median in both (raw values on **Supplementary Data 2.**). Their agreement was quantified on these co-detected genes with Spearman’s rank correlation, computed separately in two regimes split at a median STAR TPM of 5: a typical-expression regime (*≥* 5) and a very-low-expression regime (*<* 5), the latter near the detection limit and dominated by counting noise. To characterize the off-diagonal deviations, the co-detected genes were split into two groups: an off-diagonal group with a median STAR TPM *≤* 10^2^ and a median merome estimate *≥* 2 *×* 10^3^, and a diagonal group of the remaining genes. For each gene, the mean normalized sequence entropy of its 10-mers was then compared between the two groups with a Welch’s two-sided t-test.

### Detection of known biomarkers

Across the three biomarker classes, merome estimates were matched to the reference call-sets by TCGA barcode, and F1 scores were computed only over samples present in the corresponding reference (samples from projects absent from a reference contributed to detection frequencies but not to F1). For SNV hotspots the reference provided only sample-level identifiers, so samples carrying multiple aliquots were discarded to avoid ambiguous matching; for fusions and viruses the reference provided aliquot-level identifiers, so positivity was assigned unambiguously and no sample was discarded.

All HPV strains with a prevalence of more than 10 positive samples across all TCGA projects, as reported by^26^, were included: HPV16 (NC_001526.4), HPV18 (NC_001357.1), and HPV45 (J04353), and their E6 and E7 genes were used as query sequences; a sample was classified as positive if the merome estimate for E6 or E7 was non-zero, whereas a reference sample was considered positive at a minimum of 100 RPHM (reads per hundred million) ^26^. For the calculation of F1 (Fig 2c), the intersection with the merome approach was restricted to the 23 cancer types covered by the reference; the 10 uncovered TCGA projects contributed only to the frequency of positive samples. For Fig. 2a, the display was restricted to HPV16 E7 in the seven projects with at least one positive result (CESC, HNSC, SARC, BLCA, LGG, LIHC, LUSC). A minority of HPV16 merome-positive, reference-negative samples carried another documented HPV strain, consistent with inter-strain E6/E7 homology or genuine co-infection.

The hotspots^27^ were mapped to genomic coordinates using jvarkit v2024.08.25 backlocate^89^ and restricted to those with a unique mapping (*n* = 835), then filtered to retain only those hotspots with at least one TCGA project containing at least 10 positive samples according to the reference. For each selected hotspot, 25-bp query sequences were generated centered on the mutated codon and covering all synonymous codons (e.g., BRAF V600E: all codons encoding V as the reference, all those encoding E as the variant). A sample was classified as positive when its variant allele fraction, computed as the summed variant-codon estimates divided by the summed variant-plus-reference-codon estimates, exceeded 10%. Because the cBioPortal call-set^32,33^ derives from whole-exome sequencing whereas the merome measures expression, merome-positive but reference-negative calls are expected, particularly for variants expressed at low abundance.

All fusions with a prevalence of more than 10 positive samples across all TCGA projects, as reported by^28^, were included. For each fusion, 13-bp sequences flanking each side of the junction were concatenated into 26-bp queries. A sample was classified as positive if the merome estimate was non-zero for at least one queried junction. The intersection of the samples with those covered by^28^ defined the evaluation cohort for the F1 score. For the TMPRSS2::ERG comparison in prostate cancer, each TCGA-PRAD sample was cross-classified by its STAR-Fusion reference status and by the number of fusion junctions detected in the merome; reference calls supported by fewer than five junction reads were labelled as low-coverage. This distinguished samples concordant with the reference, samples in which the merome recovered additional junctions or a junction in a reference-negative sample, and reference calls attributable to low coverage or to breakpoints called differently by STAR-Fusion. The lower F1 score of FGFR3::TACC3 in Fig. 2c comes from reads that STAR-Fusion did not call as fusion-supporting, concentrated in samples profiled at a single sequencing center. The lower F1 of ABR::YWHAE comes from single-read STAR-Fusion calls, which fall below the detection threshold of our needle meromes, since each minimizer must be observed at least twice to be indexed (cutoff 2).

In the biomarker summary heatmap, a biomarker was displayed when it reached at least 10 positive samples: across all projects for oncoviruses and fusions, or within a single project for SNV hotspots. Detection proportions were colour-scaled with a square-root mapping (PowerNorm, *γ*= 0.5) and F1 scores on a [0.5, 1] scale; rows (projects) and columns (biomarkers) were ordered by hierarchical clustering (Ward linkage on Euclidean distances).

### UMAP projections

For bulk data, GTEx and TCGA meromes were queried with a single shared panel of 50,000 reads drawn at random from across the cohort; the same embedding computed over a range of panel sizes (5, 10, 50, 100, 500, 1,000, 10,000 and 50,000 reads) is shown in Fig. S3. For single-cell SMART-Seq data, per-cell matrices of reads (merome) or genes (STAR TPM) were used with 10,000 features, restricted to biological replicate 1. Expression values were log1p-transformed without further scaling, reduced by PCA (50 components, full SVD solver) and embedded with UMAP (scanpy v1.10.4^83^, umap-learn v0.5.7^90^; n_neighbors = 25, min_dist = 0.5, spread = 1.0, random_state = 42). For the smaller panels, the number of principal components and the neighbourhood size were capped at the data dimensions (n_components = minimum of 50, n_features *−* 1 and n_samples *−* 1; n_neighbors = minimum of 25 and n_samples *−* 1); the separate principal-component projections were instead computed on features scaled to a maximum value of 10.

### Supervised classification

For each cohort, a pool of one million reads was randomly sampled (without replacement) from a subset of samples: 330 TCGA samples (10 per project across 33 projects); 9,225 GTEx samples across 30 tissues; and the union of these two pools for TCGA+GTEx (equal contribution: 500k reads for GTEx and 500k reads for TCGA). For SMART-Seq thymocytes, the draw was stratified across the 1,489 preserved cells by thymic subpopulation (7 subpopulations, also totaling one million reads).

Four configurations were tested, each pairing a read pool with its target merome: GTEx, TCGA, joint TCGA+GTEx, and SMART-Seq thymus single-cell. For the TCGA and joint TCGA+GTEx configurations, non-primary samples were excluded. A STAR TPM reference (gene-level, or single-cell for SMART-Seq) was compared to each configuration. The number of features varied across 11 values from 1 to 50,000 (1, 5, 10, 20, 30, 50, 100, 500, 1,000, 10,000, 50,000; 50,000 corresponds to the order of magnitude of the number of annotated genes), with 100 stratified runs per combination.

The expression values (merome estimates or TPM STAR) were transformed using log1p, then split into training and test sets according to an 80%/20% split stratified by label (GTEx tissue, TCGA project, or thymic subpopulation, depending on the configuration). For each run, a Random Forest classifier ^91^ (sklearn.ensemble. RandomForestClassifier, scikit-learn v1.5.2^92^, n_estimators=300, max_leaf_nodes=15,000, random_state=0, criterion=entropy) was trained on the training split without cross-validation and then applied to the test split. The importance of each feature for classification was determined using the entropy-based importance score (mean decrease in impurity) from the Random Forest classifier. The difference confusion matrix was computed between the row-normalised merome and STAR-TPM confusion matrices of the SMART-Seq thymocytes and oriented so that positive values always favour the merome: on the diagonal, the gain in correctly classified cells (merome *−* STAR); off the diagonal, the reduction in misclassifications (STAR *—* merome). Classification performance was summarized by the macro-averaged F1 score, accuracy, precision and recall (scikit-learn ^92^); the metric-versus-feature-count curves report the mean *±* standard deviation over the 100 runs (Fig. 3e,f), and the thymocyte panels the corresponding per-run distribution at 10,000 features (Fig. 4d,e). Displayed confusion matrices were row-normalized and evaluated at 1,000 features for the bulk cohorts (TCGA, GTEx and joint) and at 10,000 features for the single-cell thymocytes, aggregating the 100 runs. In the per-sample prediction heatmaps (50 features), each sample’s predictions were tallied over the 100 runs and samples were ordered by hierarchical clustering (Ward linkage, Euclidean distance) of their prediction-frequency profiles.

To characterize the genomic origin of the discriminative reads, feature importances (mean decrease in impurity) from the joint GTEx+TCGA classifier (50,000 features, 50 runs) were pooled across runs and renormalized. Each read was assigned both to a single HOMER region category (3’UTR, CDS exon, non-coding, 5’UTR, TTS, intron, promoter-TSS or intergenic) and to its alignment signature (perfect, SNV, spliced, insertion or deletion). The informational enrichment of a category was its importance-weighted proportion of reads divided by its raw proportion, a value of one indicating no enrichment; categories supported by fewer than five reads were discarded.

### Alignment-last

The reads were aligned using STAR v2.7.11b^22^ in twopassMode Basic against GRCh38.p14 and Gencode v45 (human) or against mm10 and Gencode vM22 (mouse), with the following parameters: –outFilterMultimapNma× 1000, –winAnchorMultimapNma× 1000, –outSAMunmapped Within, –outSAMattributes NH HI NM MD AS nM jM jI, –chimOutType SeparateSAMold, –bamRemoveDuplicatesType UniqueIdentical, – outFilterMismatchNoverLmax 0.04, –outMultimapperOrder Random, –chimSegmentMin 10, –chimJunctionOverhangMin 10, and –limitOutSAMoneReadBytes 1000000. The main and chimeric alignment files produced by STAR were concatenated using samtools merge (samtools v1.9), converted to BAM (samtools view), and then to BED using bedtools bamtobed -split (bedtools v2.31.1). Each read was then annotated by HOMER annotatePeaks.pl (HOMER v5.1)^23^ on the hg38 or mm10 genome, simultaneously providing gene annotations (gene name, genomic region, proximity to features) and transposable element annotations (class, family, and subfamily). Each read was also assigned a taxonomic origin with Kraken2 v2.1.3^68^ (standard database k2_standard_20240904). Each read was given an alignment signature derived from its CIGAR string and edit distance: perfect (only matches, NM = 0), or otherwise SNV, spliced, insertion or deletion. These signatures defined the read subsets used in the differential analyses below.

### Differentially expressed reads

Read-level differential expression used a single run, shared by the lncRNA, transposable-element and unmapped-read analyses. One million reads stratified by project (500,000 in GTEx and 500,000 in TCGA, the latter restricted to primary tumours) were queried against the GTEx and TCGA meromes. Each project was contrasted against all remaining samples (other tumours and healthy tissues) with pydeseq2 v0.5.3 (DESeq2^93^) by binarizing the project label into target versus rest, with rest as the reference level, under a Wald test (*α* = 0.05) with Cook’s-distance refitting. To bound memory, the reads were shuffled (random seed 42) and split into independent batches of 1,000 reads, each fitted by a separate DESeq2 run with size factors estimated within the batch; the resulting P values were pooled and the Benjamini–Hochberg correction recomputed globally per condition. Differentially expressed reads were used directly from this run for the transposable-element analysis; the restriction to the ‘perfect’ alignment signature (a CIGAR of matches only, NM = 0) was applied solely to the lncRNA candidate selection described next.

Tumour-overexpressed lncRNA reads were selected from this differential analysis across the 11 TCGA projects with a CPTAC equivalent (BRCA, COAD, GBM, HNSC, KIRC, LUAD, LUSC, OV, PAAD, READ and UCEC). Reads shorter than 25 nt were discarded, and a read was treated as non-coding only when annotated ‘non-coding’ by HOMER and never annotated as protein-coding across its occurrences. Within each project, such reads with fold change *≥* 4 and adjusted *P ≤* 5 *×* 10*^−^*^3^ were kept, and the 100 top-ranked by adjusted P and the 100 top-ranked by log2 fold change were taken and unioned. The retained reads were deduplicated by sequence and restricted to the ‘perfect’ alignment signature, yielding 391 lncRNA reads (provided in **Supplementary Data 3**).

Transposable-element reads were taken from the same differential analysis, but without the candidate-selection or ‘perfect’-signature restriction applied to lncRNAs: within each tumour, reads with adjusted *P ≤* 0.05 carrying a genuine HOMER transposable-element annotation were ranked by descending log2 fold change and the top 100 retained, with their class, family and subfamily taken from HOMER (**Supplementary Data 4** for ACC and **Supplementary Data 5** for SARC). LTR10D in ACC and L1ME2z in SARC are shown as illustrative examples. For genes highlighted in the volcano plots, the number of differentially expressed reads annotated by HOMER to the gene and falling in the up-regulated region (log2 fold change *≥* 1, adjusted *P ≤* 0.05) was counted, for example 650 reads for miR-483 in ACC and 22 reads for CAPN6 in SARC.

The 391 lncRNA reads were embedded by UMAP, as above, independently per cohort (TCGA and GTEx), GTEx sub-tissues being collapsed to their parent tissue. The four reads exceeding log2 fold change 4 between TCGA and GTEx selected with a Wilcoxon rank-sum test (scanpy, reference GTEx) and deduplicated to unique gene symbols, were displayed as a per-read min–max-scaled (0–1) heatmap across the GTEx, TCGA and CPTAC cohorts, samples being ordered within each cohort by average-linkage hierarchical clustering (Euclidean distance) of their per-project means. Per-read detection was summarized by scatter plots in which each point is a project (TCGA and CPTAC tumours, GTEx tissues), the x axis the fraction of positive samples and the y axis the mean estimate over positive samples (symlog scale, linear threshold 1); a read was considered near-silent in a tissue when detected in fewer than 5% of its samples.

Differential expression for the single-cell exhaustion dataset used a separate pydeseq2 run contrasting Tpex and Tex cells (10x, mouse), the reads annotated by the mouse alignment-last pipeline (mm10, Gencode vM22). Aligned reads with a recorded sequence, without the ‘perfect’ restriction, passing |log2 fold change| *>* 1 and adjusted *P* < 0.05 were retained; genes and transposable-element subfamilies were then ranked by the mean log2 fold change of their differentially expressed reads (features with at least five such reads), reads lacking a transposable-element annotation being excluded from the TE ranking.

### Merome-based positivity and expression comparisons

A sample was called positive for a read or sequence when its depth-normalized merome estimate exceeded zero. For per-read expression comparisons boxplots, the estimates of the positive samples in the focus tumour were compared with those of, respectively, the other tumours and the healthy tissues using two-sided Mann–Whitney U tests (SciPy^84^). Overlaid jitter points were sub-sampled for legibility (all focus-tumour samples, 10% of other tumours and 5% of healthy tissues, fixed seed), and non-positive estimates were omitted on the logarithmic axis.

### Survival analysis

Overall survival was analysed with Kaplan–Meier estimates and two-sided log-rank tests (lifelines v0.30.1^94^), using overall-survival time and status from the TCGA Pan-Cancer clinical resource^95^, matched to the samples through the 12-character patient barcode, so that a patient contributing several sequencing files shares a single survival record across them. Within the focus tumour, read-positive (estimate *>* 0) and read-negative samples were compared. For sarcoma, the comparison was repeated within each histological subtype represented by at least 15 samples, with the remaining subtypes pooled into an ‘other’ group.

### Association with molecular and histological subtypes

Associations between read positivity and subtype were tested on contingency tables with Pearson’s *χ*^2^ test of independence, or Fisher’s exact test for 2 *×* 2 tables, with Cramér’s V as effect size (SciPy). For each table, adjusted standardized residuals (Agresti) flagged enriched (*z* > 0) or depleted (*z* < 0) categories from two-sided z-derived P values with Bonferroni correction (*α* divided by the number of rows). The ACC molecular subtypes (cluster-of-clusters, steroid phenotype and C1A/C1B^96^) were taken from the TCGA-ACC reference classification^61^ and matched to samples through the 12-character patient barcode; the sarcoma histological subtype was the sample’s primary diagnosis truncated at its first qualifier (e.g. ‘Leiomyosarcoma, NOS’ *→* ‘Leiomyosarcoma’), subtypes with fewer than 15 samples being pooled into an ‘other’ group. For ubiquitously expressed elements such as L1ME2z, the prevalence per subtype was reported with 95% Wilson confidence intervals.

### Read-versus-full-element analysis

To test whether a single read faithfully reports its element, the full-length sequence of the element was retrieved with samtools faidx at its RepeatMasker coordinates (release 4.3.1, GRCh38), added as an additional merome query, and the positivity, survival and subtype analyses were repeated (Fig. S8, Fig. S10). The L1ME2z read spans two adjacent RepeatMasker elements, AluSz and L1ME2z; the chimaeric sequence and each element were queried separately to attribute the signal to the L1ME2z component (Fig. S10).

### Unaligned reads and chimeric junctions

This analysis was run on CPTAC rather than TCGA, whose tumour taxonomic signal has been shown to be dominated by contamination and analysis artefacts^67^. One million reads stratified by CPTAC project were queried against the CPTAC meromes and analysed by read-level differential expression as above (pydeseq2, each project contrasted one-vs-rest, Wald test *α* = 0.05 with Cook’s-distance refitting). Reads left unaligned by STAR (SAM flag 0×4) were retained and assigned a taxonomic origin with Kraken2 (k2_standard_20240904) as described above.

For each project, unaligned reads were ranked by their DESeq2 Wald statistic and tested for enrichment by taxonomic origin with preranked gene-set enrichment analysis (gseapy v1.1.12 prerank; 1,000 permutations, minimum set size 20, seed 42). Reads whose Kraken2 assignment fell under the NCBI ‘cellular organisms’ node (taxon 131567) were pooled into a single set (8,009 reads) and reads assigned to the human genome (taxon 9606) into another (10,682 reads).

Human unaligned reads enriched in serous ovarian carcinoma were screened for chimeric junctions with a dedicated STAR v2.7.11b pass using the parameters recommended by Arriba^39^ (–chimOutType Junctions, –chimSegmentMin 10, –chimJunctionOverhangMin 10, –chimScoreDropMax 30, –chimScoreJunctionNonGTAG 0, –chimScoreSeparation 1, – himSegmentReadGapMax 3, –chimMultimapNmax 50, –alignSJstitchMismatchNmax 5-1 5 5, –peOverlapNbasesMin 10, –alignSplicedMateMapLminOverLmate 0.5, with read-filtering thresholds relaxed). In the OV volcano, unaligned human reads whose read name appeared in this chimeric output were highlighted; the dashed lines mark the significance thresholds, an adjusted P of 0.05 and an absolute log2 fold change of 2.

For each chimeric junction, a 26-bp query (13 bp on each side of the breakpoint) was reconstructed from GRCh38.p14 with samtools faidx, with strand handling. Junctions were discarded when the breakpoint fell in a repeat or microhomology, lay on a contig absent from the reference (e.g. viral contigs), carried a non-templated insertion, was not spanned by a split read, or could not reach 26 bp (2,390 of 6,677 junctions retained). These queries were searched against the CPTAC-OV, TCGA-OV and GTEx ovary meromes; junctions positive in at least one sample of any cohort were kept (1,332). For each cohort, the fraction of positive samples and the mean estimate among positive samples were summarized by bootstrap (10 resamples; point, mean; error, *±*2 s.d.).

## Data availability

The TCGA RNA-seq data analyzed in this study are available via the NCI Genomic Data Commons Data Portal (https://portal.gdc.cancer.gov/) under dbGaP accession number phs000178. GTEx v8 RNA-seq data are available via the AnVIL platform (https://gen3.theanvil.io/) under dbGaP accession phs000424. CPTAC RNA-seq data are available via the NCI Genomic Data Commons under dbGaP accession phs001287. The SMART-Seq scRNA-seq dataset of mouse thymocytes^46^ and the 10x scRNA-seq dataset of CD8^+^ TILs^21^ are available under the accession numbers indicated in the original publications. Reference datasets used as ground truth: pan-cancer SNV hotspots at https://www.cancerhotspots.org/^27^; SNV call-sets at https://www.cbioportal.org/^32,33^; TCGA pan-cancer fusion landscape^28^ (PMC5916809); TCGA viral reference^26^ (PMC4919655). HPV reference sequences: NCBI accessions NC_001526.4 (HPV16), NC_001357.1 (HPV18), J04353 (HPV45).

## Code availability

Merome Explorer, a web platform enabling the scientific community to query RNA sequences of interest and obtain quantitative expression estimates across tens of thousands of samples within minutes, is available online (the platform URL and access credentials are withheld in this preprint version). The readdiff command-line tool, which implements the alignment-last pipeline to identify and explore differentially expressed reads, will be released on GitHub and is currently available as a zip file attached to submission.

## Acknowledgments

We thank Rayan Chikhi, Daniel Gautheret and Knut Reinert for helpful discussions and feedback on this work.

## Supplementary Datas

**Supplementary Data 1.** Per-biomarker detection frequencies and F1 scores against the reference call-sets across the 33 TCGA projects (oncoviruses, SNV hotspots and gene fusions).

**Supplementary Data 2.** Median STAR TPM and median merome estimate per gene, for the co-detected Gencode v45 genes.

**Supplementary Data 3.** The 391 tumour-overexpressed lncRNA reads, with sequence, genomic annotation and per-project differential statistics.

**Supplementary Data 4.** Top-100 transposable-element reads upregulated in adrenocortical carcinoma (TCGA-ACC).

**Supplementary Data 5.** Top-100 transposable-element reads upregulated in sarcoma (TCGA-SARC).

## Extended Data Figures

**Fig. S1.**
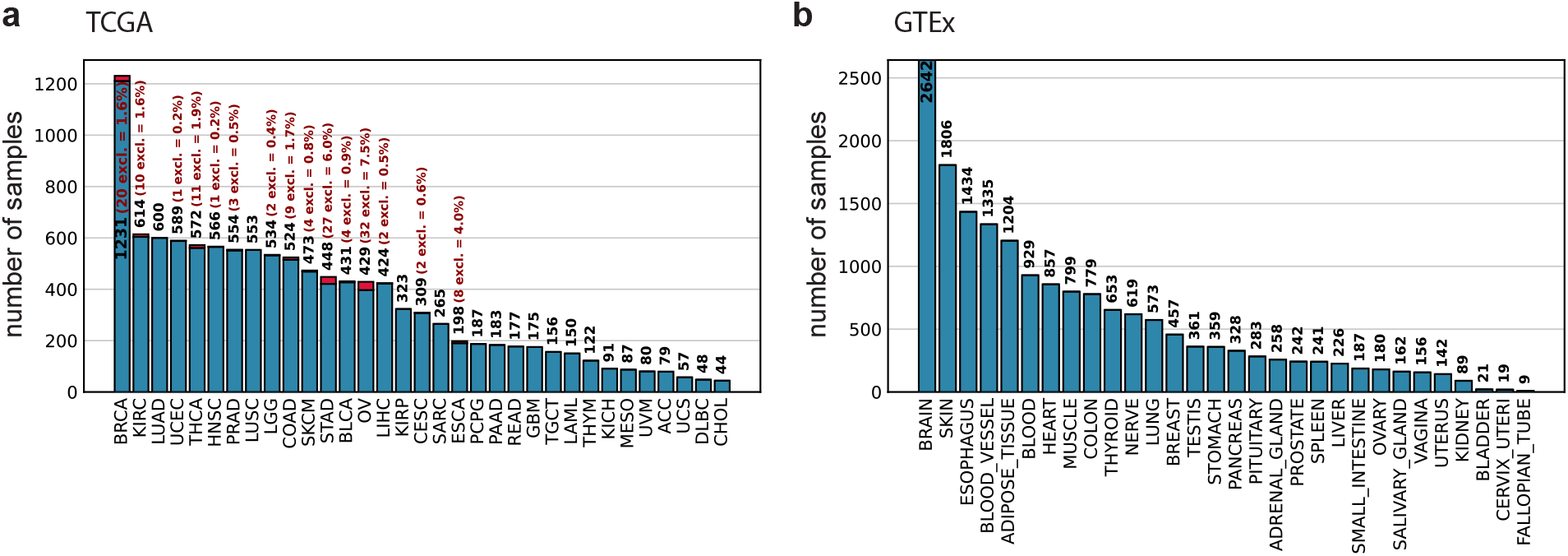
Samples selected for the construction of the TCGA and GTEx meromes. **a**, Samples included vs. excluded during the construction of TCGA meromes, by tumor project (*n* = 33). A sample is excluded when the size of its fastq.gz file exceeds the project threshold (13 GB by default; 23 GB for ESCA, GBM, LUAD, LUSC, OV, STAD), a criterion determined by the memory available for merome construction. Total: 11,137 samples included out of 11,273 eligible (98.79%); 15 out of 33 projects had at least one exclusion. **b**, Samples selected by GTEx tissue type (*n* = 30 main tissues divided into 54 tissue subdivisions). All 17,350 eligible samples were included (100%).

**Fig. S2.**
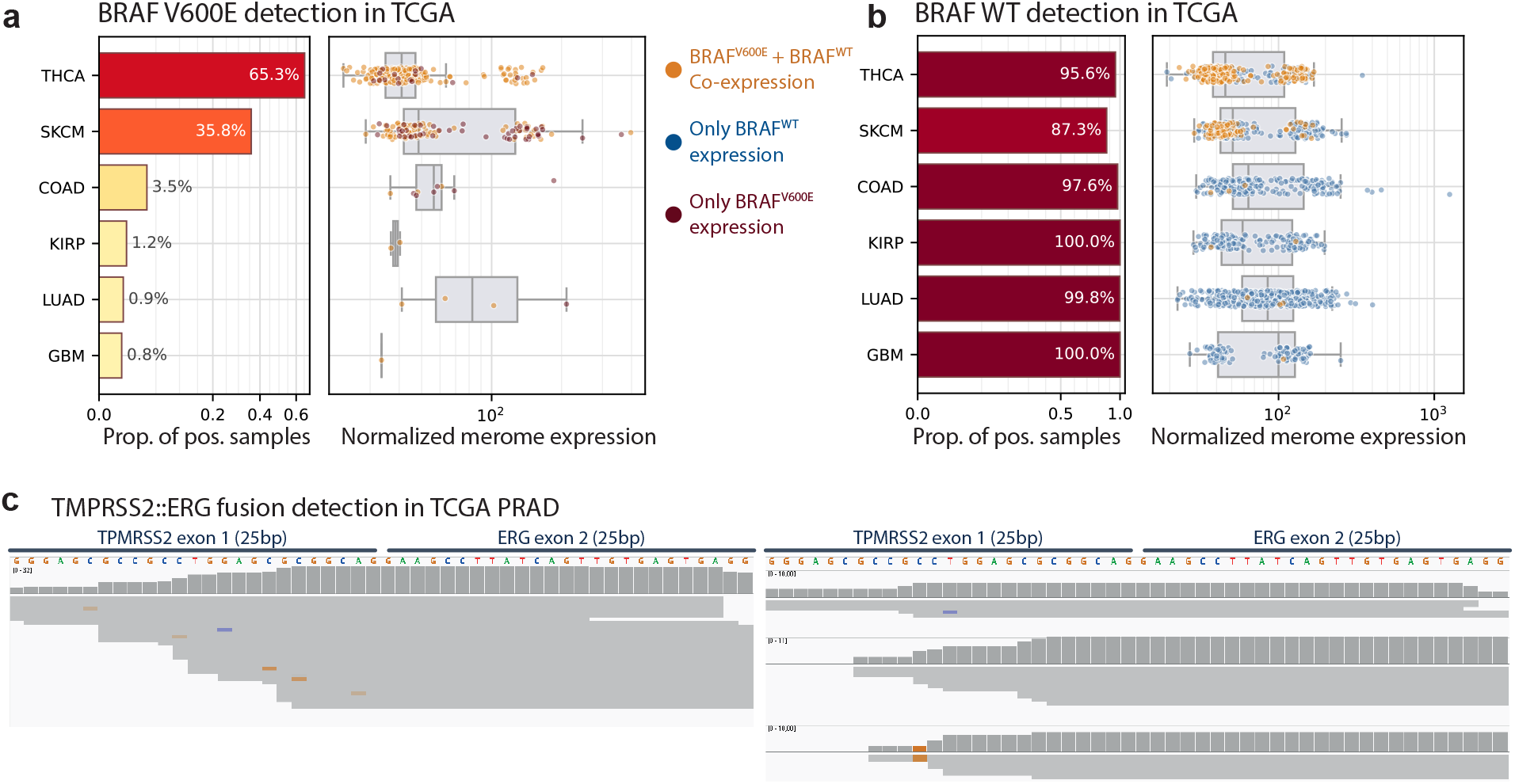
Merome detection of the BRAF V600E variant and the TMPRSS2-ERG fusion. **a**, Detection of the BRAF V600E variant in six TCGA projects. Left: proportion of samples positive for the variant (THCA 65.3%, SKCM 35.8%, COAD 3.5%, KIRP 1.2%, LUAD 0.9%, GBM 0.8%). Right: Normalized merome expression among positive samples, with each point colored according to allelic expression pattern (orange: co-expression of BRAF V600E and BRAF WT; dark red: BRAF V600E alone; blue: BRAF WT alone). **b**, Same plot for the detection of the wild-type allele associated with BRAF V600E in projects that expressed the mutation. The wild-type allele is expressed in nearly all samples from the relevant projects (87.3–100%). **c**, Reads supporting the TMPRSS2::ERG fusion in two TCGA-PRAD samples that are positive (Fig. 2d) for the merome but negative according to the STAR-Fusion reference ^28^.

**Fig. S3.**
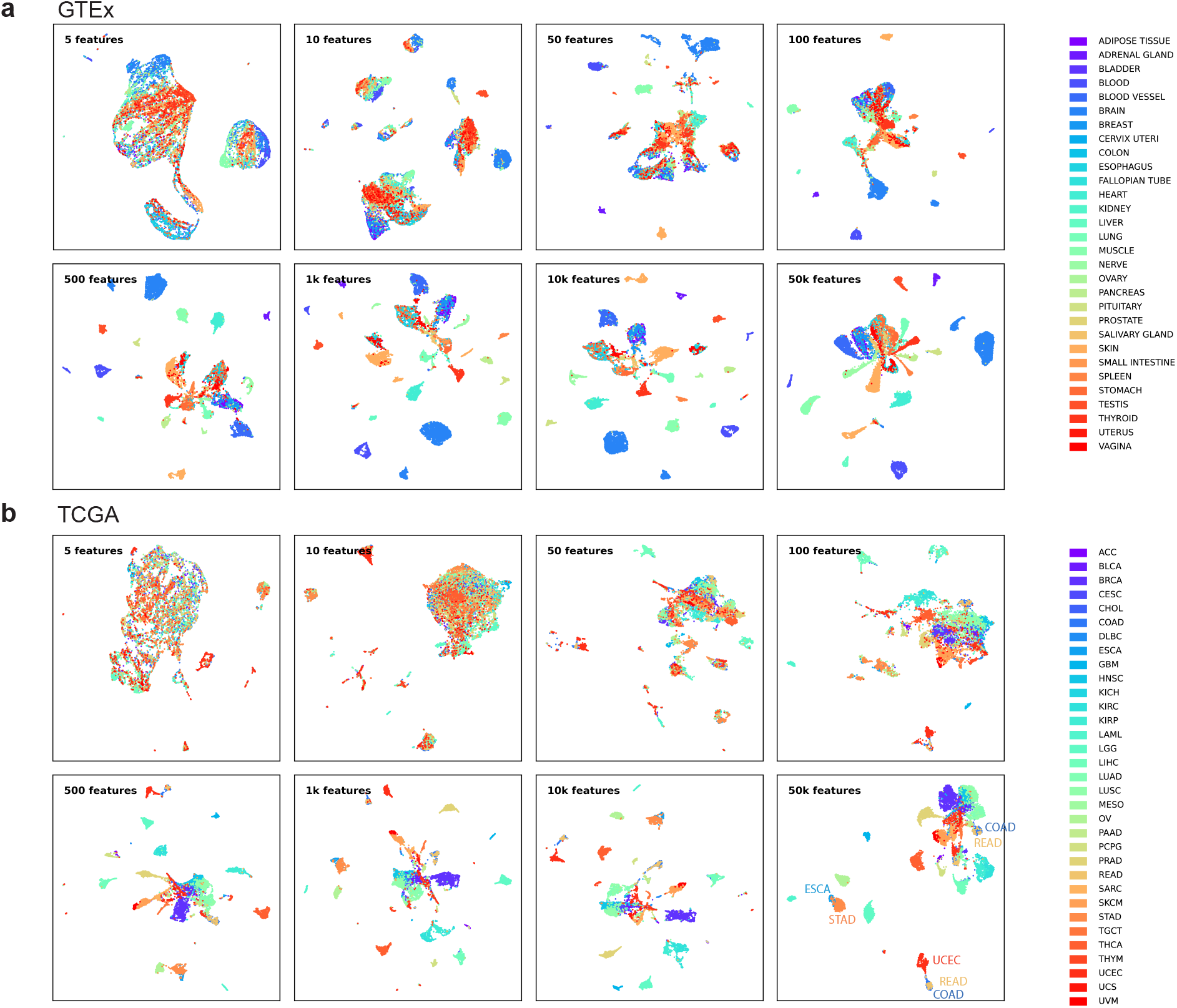
Cohort separation improves with the number of randomly sampled reads. **a,** UMAP projections of GTEx samples based on 5, 10, 50, 100, 500, 1,000, 10,000, and 50,000 randomly sampled reads; each point represents a sample, colored according to its tissue type (30 tissues). **b,** Same plot for TCGA samples, colored according to their tumor project (33 projects).

**Fig. S4.**
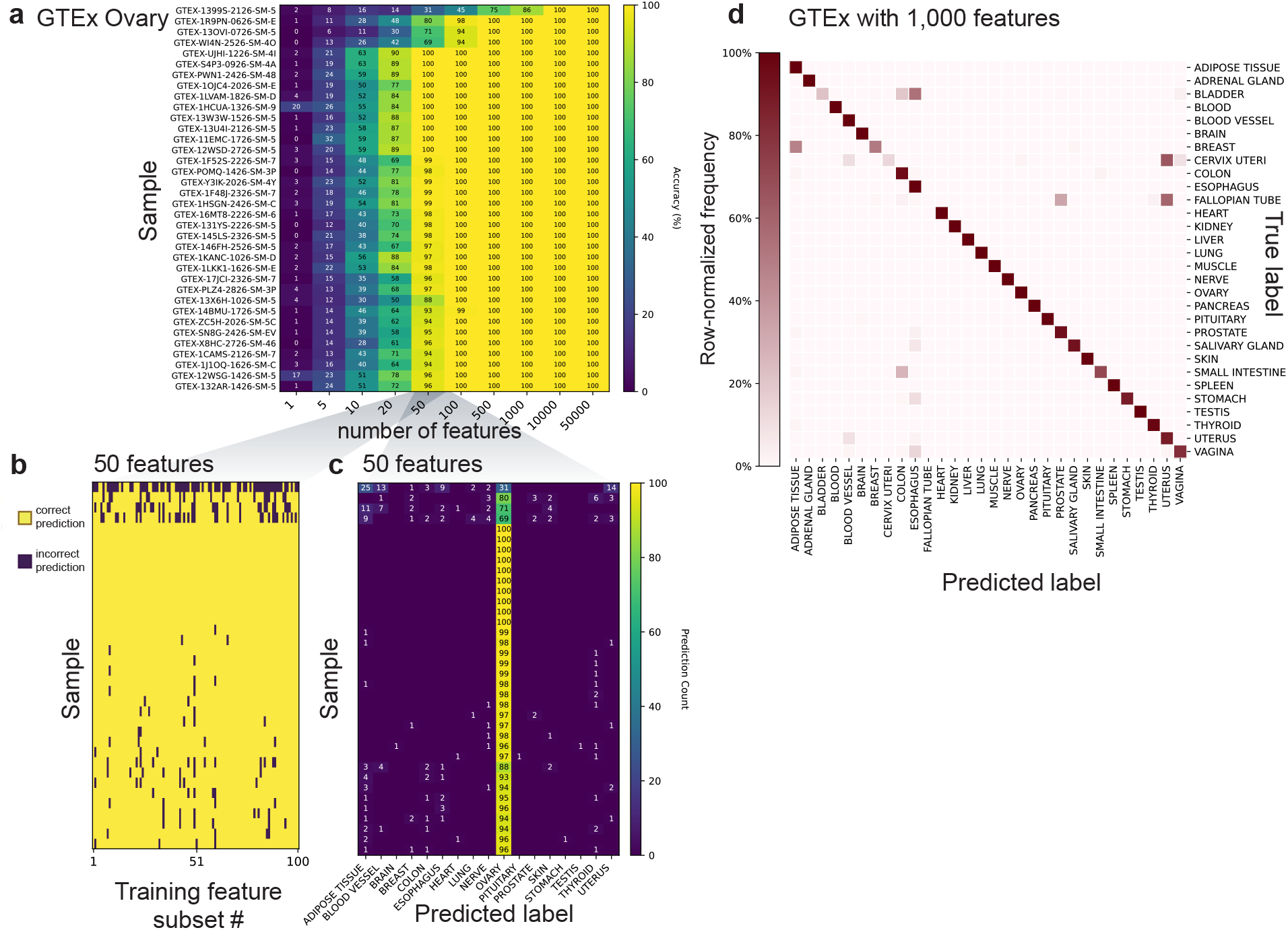
The reads accurately classify healthy GTEx tissues. **a**, Tumor classification details for GTEx Ovary true-labeled samples (GTEX-OVARY; 36 test samples) based on the number of features. Each row corresponds to a sample, and each column corresponds to a feature subset size (randomly drawn reads; 1 to 50,000); color indicates the proportion of correct predictions out of 100 independent draws. The samples are ordered by hierarchical clustering. **b**, For 50 features, prediction accuracy per sample (rows) and per feature selection (columns; 100 selections) on the GTEx Ovary test set; yellow = correct prediction, purple = incorrect prediction. **c**, For 50 features, distribution of predicted labels (columns) for each sample in the GTEx Ovary test set (rows); the color indicates the number of draws (out of 100) that assigned each label. **d**, Confusion matrix for the GTEx classifier (reads, 1,000 features; averaged over 100 runs); rows = true label, columns = predicted label, color = row-normalized frequency (30 projects, overall accuracy 96.9%).

**Fig. S5.**
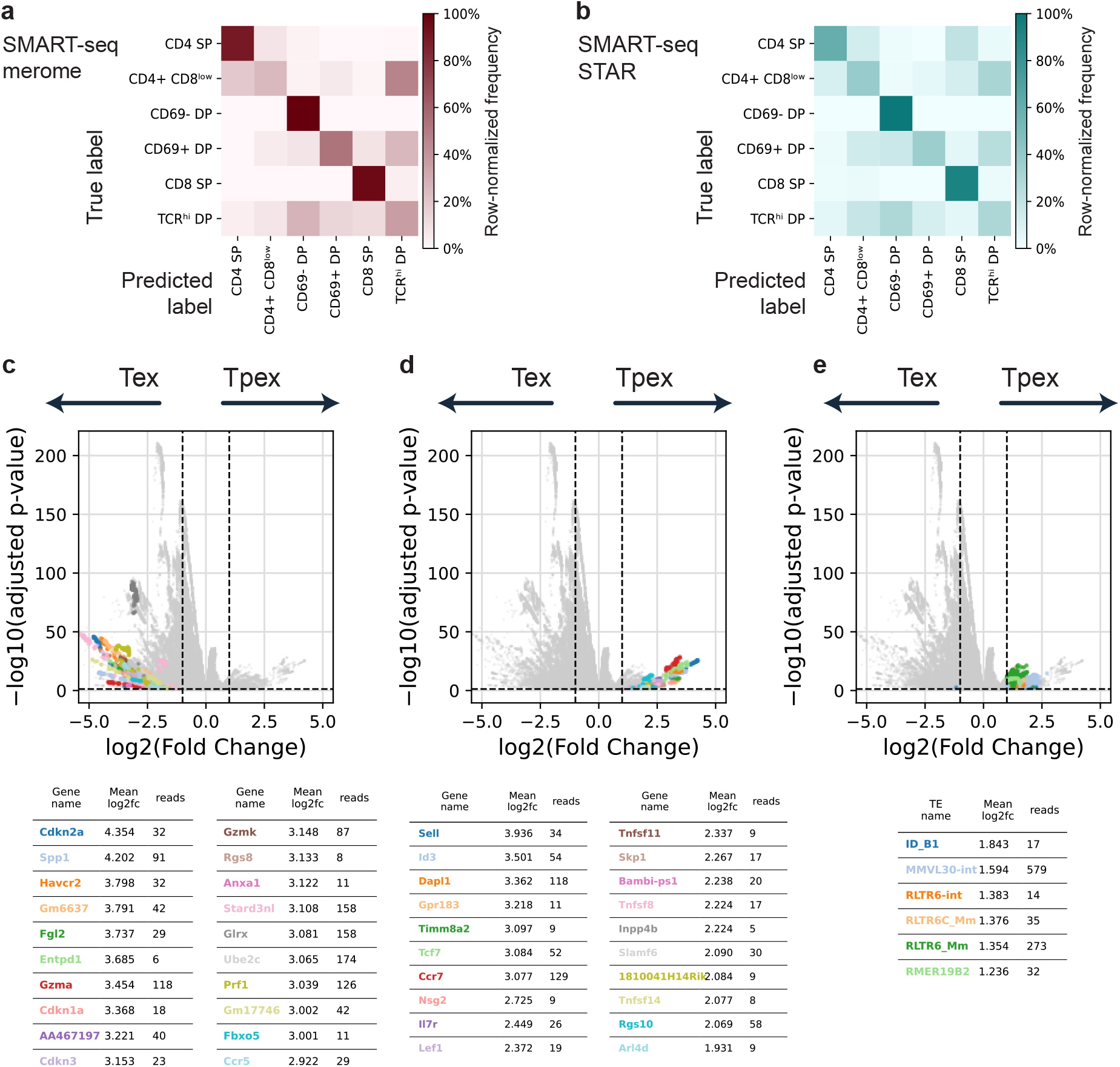
Single-cell complements: thymocyte classification (SMART-Seq) and differentially expressed reads between Tpex and Tex (10x). **a,** Confusion matrix for the Random Forest classifier (10,000 features, 100 runs) on SMART-Seq thymocytes based on the merome (rows = true class, columns = predicted class, frequency normalized by row; overall accuracy 67.0%). **b,** Same confusion matrix based on STAR-TPM gene quantification (accuracy 60.2%). **c,** Volcano plot of reads differentially expressed between Tpex and Tex; the 20 most highly overexpressed genes on the Tex side (gene names according to HOMER annotation) are highlighted and detailed in the table below (gene, score, number of reads); thresholds |log2FC| = 1 and adjusted p-value = 0.05. **d,** Same plot, showing the 20 most highly overexpressed genes on the Tpex side.

**Fig. S6.**
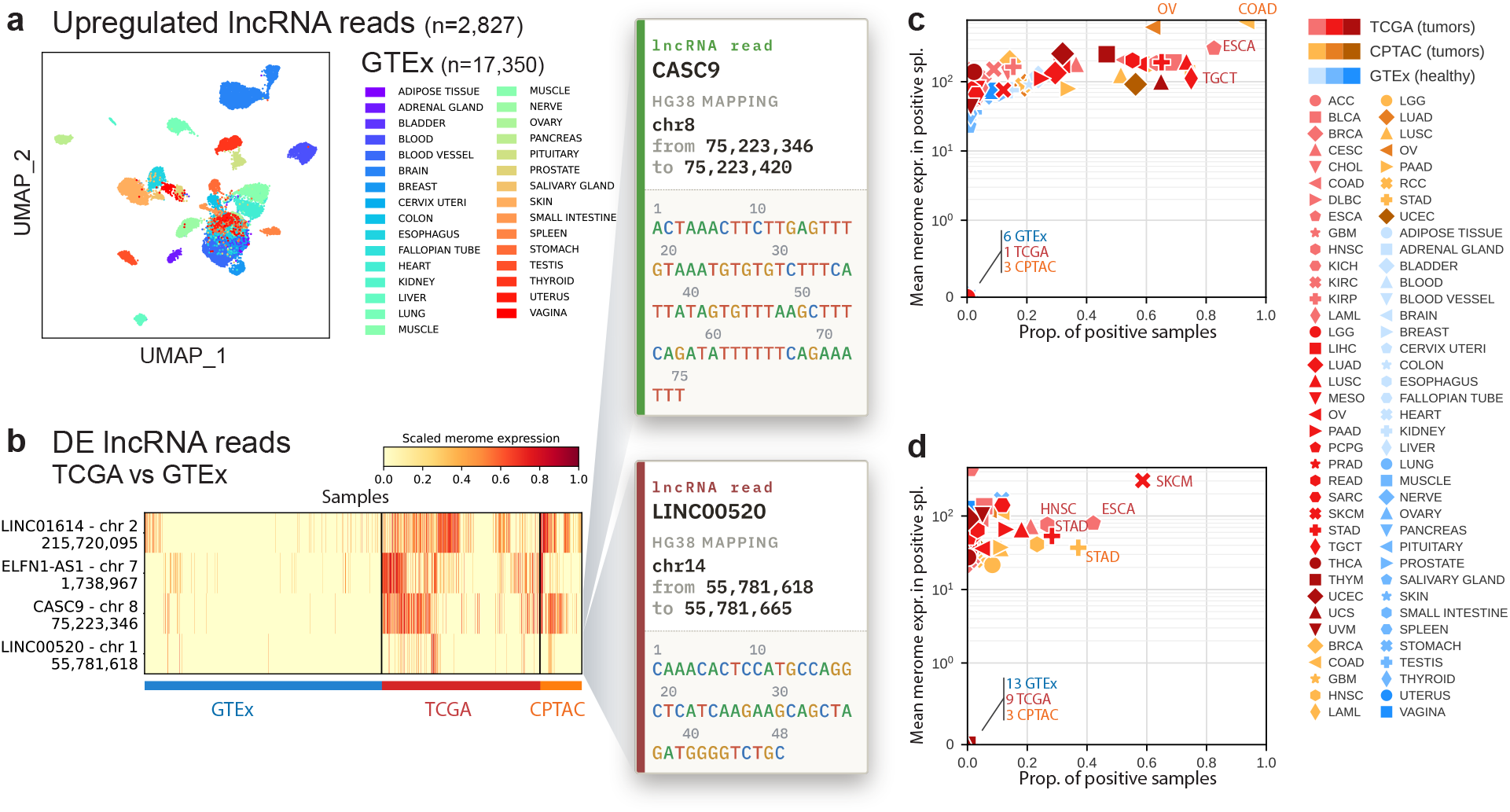
The tumor lncRNA signal is also present in healthy GTEx tissues; prevalence of CASC9 and LINC00520. **a**, UMAP of the 17,350 GTEx healthy-tissue samples (30 tissue types) from the same 391 tumor-overexpressed lncRNA reads as in Fig. 5a; each point is a sample, colored by tissue type. **b**, CASC9 read (hg38 chr8:75,223,346–75,223,420): proportion of positive samples (x-axis) versus mean merome expression among them (y-axis, symlog); each point is a project (TCGA and CPTAC tumors; GTEx healthy tissues). Detected in fewer than 5% of samples in 23 of 30 GTEx tissues. **c**, LINC00520 read (hg38 chr14:55,781,618–55,781,665): as in **b**; detected in fewer than 5% of samples in 29 of 30 GTEx tissues.

**Fig. S7.**
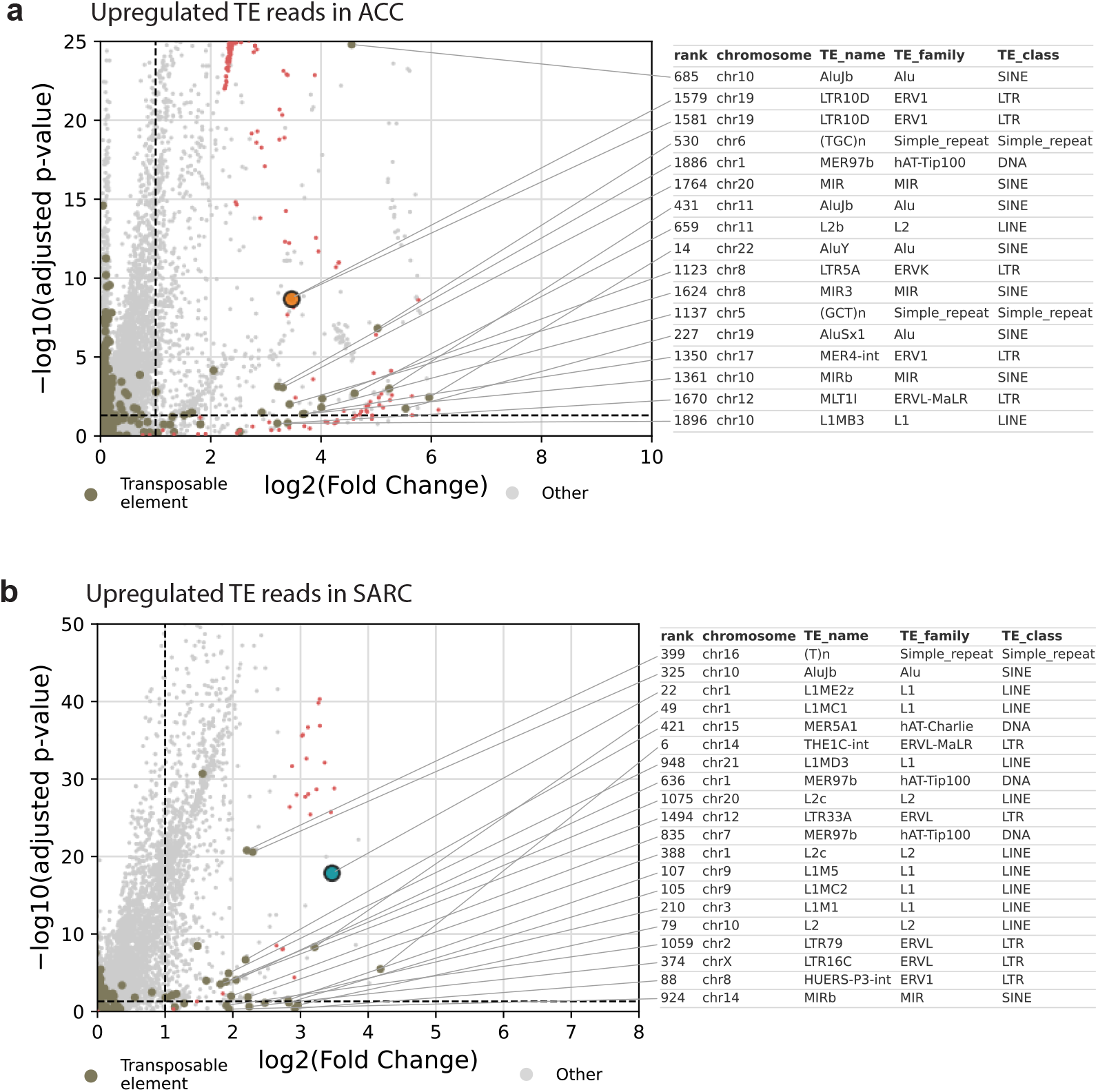
Transposable-element reads overexpressed in adrenocortical carcinoma (ACC) and sarcoma (SARC). **a**, Upregulated TE reads in ACC (TCGA-ACC versus other TCGA tumors and GTEx healthy tissues): TE reads brown, other reads gray, LTR10D dup158 orange, reads aligned to miR-483 red; thresholds |log2FC| = 1 and adjusted *p* = 0.05 (dotted lines). The table lists the 20 most overexpressed TE reads (rank, chromosome, name, family, TE class); top 100 in **Supplementary Data 4**. **b**, Same in SARC (TCGA-SARC versus the rest); L1ME2z dup127 turquoise, reads aligned to CAPN6 red; top 100 in **Supplementary Data 5**.

**Fig. S8.**
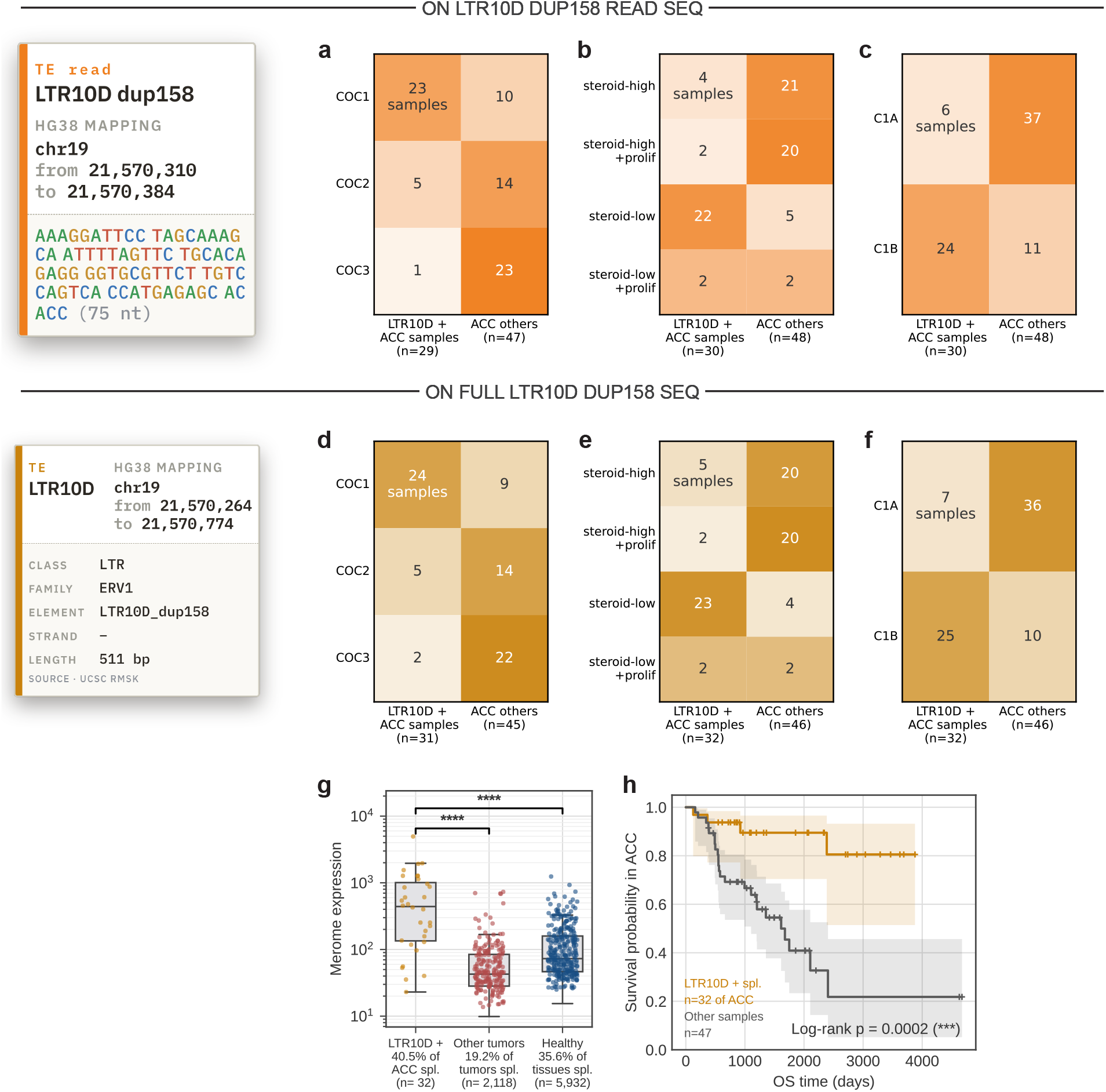
The LTR10D read identifies a well-defined ACC molecular subgroup (COC1, low-steroid phenotype, C1B), confirmed by the full-length element. Contingency tables showing LTR10D^+^ status (merome expression *>* 0) versus other ACCs, crossed with the TCGA-ACC reference classification ^61^ (color, row percentage). **a–c**, LTR10D dup158 read (chr19:21,570,310–21,570,384, 75 bp); **d–h**, full LTR10D sequence (LTR/ERV1, chr19:21,570,264–21,570,774, 511 bp). **a**, “Cluster-of-clusters” (COC) subtypes: enriched for COC1, depleted for COC3 (*χ*^2^ *p* = 1.5 *×* 10*^−^*^6^; Cramér’s *V* = 0.59; *z* = +4.96 and *−*4.14, Bonferroni-significant). **b**, Steroid phenotype (mRNA_K4): enriched in steroid-low (*z* = +5.68), depleted in high-steroidogenesis or proliferative phenotypes (*χ*^2^ *p* = 1.4 *×* 10*^−^*^7^; *V* = 0.67). **c**, C1A/C1B subtypes: enriched in C1B, depleted in C1A (Fisher’s exact test, *p* = 1.3 *×* 10*^−^*^6^; *V* = 0.53). **d–f**, Same for the full LTR10D sequence, identical enrichments (COC *p* = 2.2 *×* 10*^−^*^6^; steroid *p* = 8.5 *×* 10*^−^*^8^; C1A/C1B *p* = 8.8 *×* 10*^−^*^7^). **g**, Full-sequence expression: 40.5% of ACC positive (*n* = 32) versus 19.2% of other tumors (*n* = 2, 118) and 35.6% of healthy tissues (*n* = 5, 932) (Mann–Whitney). **h**, Overall survival in ACC, LTR10D^+^ (full sequence) versus others: better in LTR10D^+^ (log-rank *p* = 0.0002). https://figures.qwann.fr/ S10.pdf

**Fig. S9.**
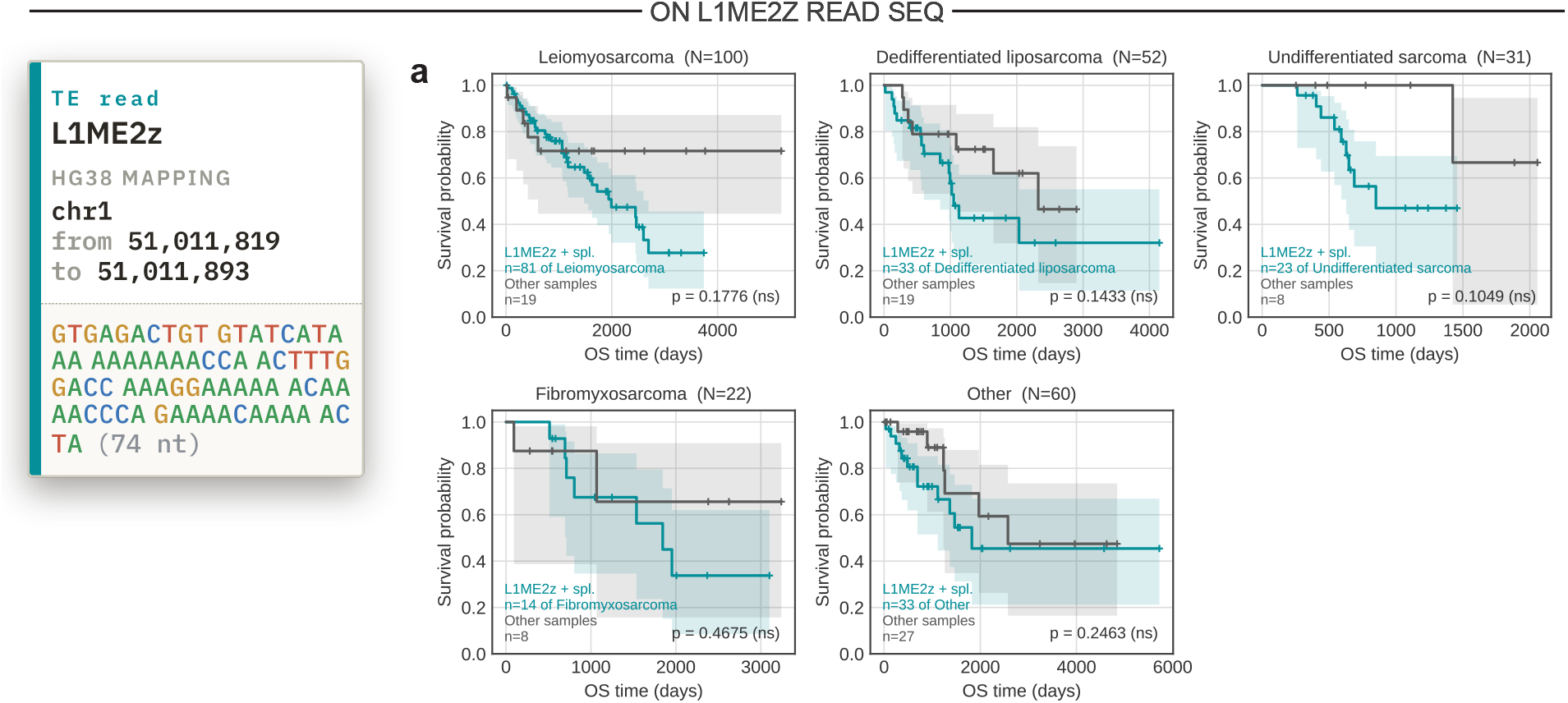
Survival associated with the L1ME2z read, by sarcoma histological subtype. **a**, Overall survival (Kaplan–Meier) in L1ME2z^+^ (turquoise) versus L1ME2z*^−^* (gray) sarcomas, within each subtype with at least 15 samples: leiomyosarcoma (*N* = 100), undifferentiated liposarcoma (*N* = 52), undifferentiated sarcoma (*N* = 31), fibromyxosarcoma (*N* = 22) and other subtypes (*N* = 60). No comparison is significant (log-rank *p* = 0.18, 0.14, 0.10, 0.47, 0.25; subtype samples too small); curve order mirrors Fig. 6g.

**Fig. S10.**
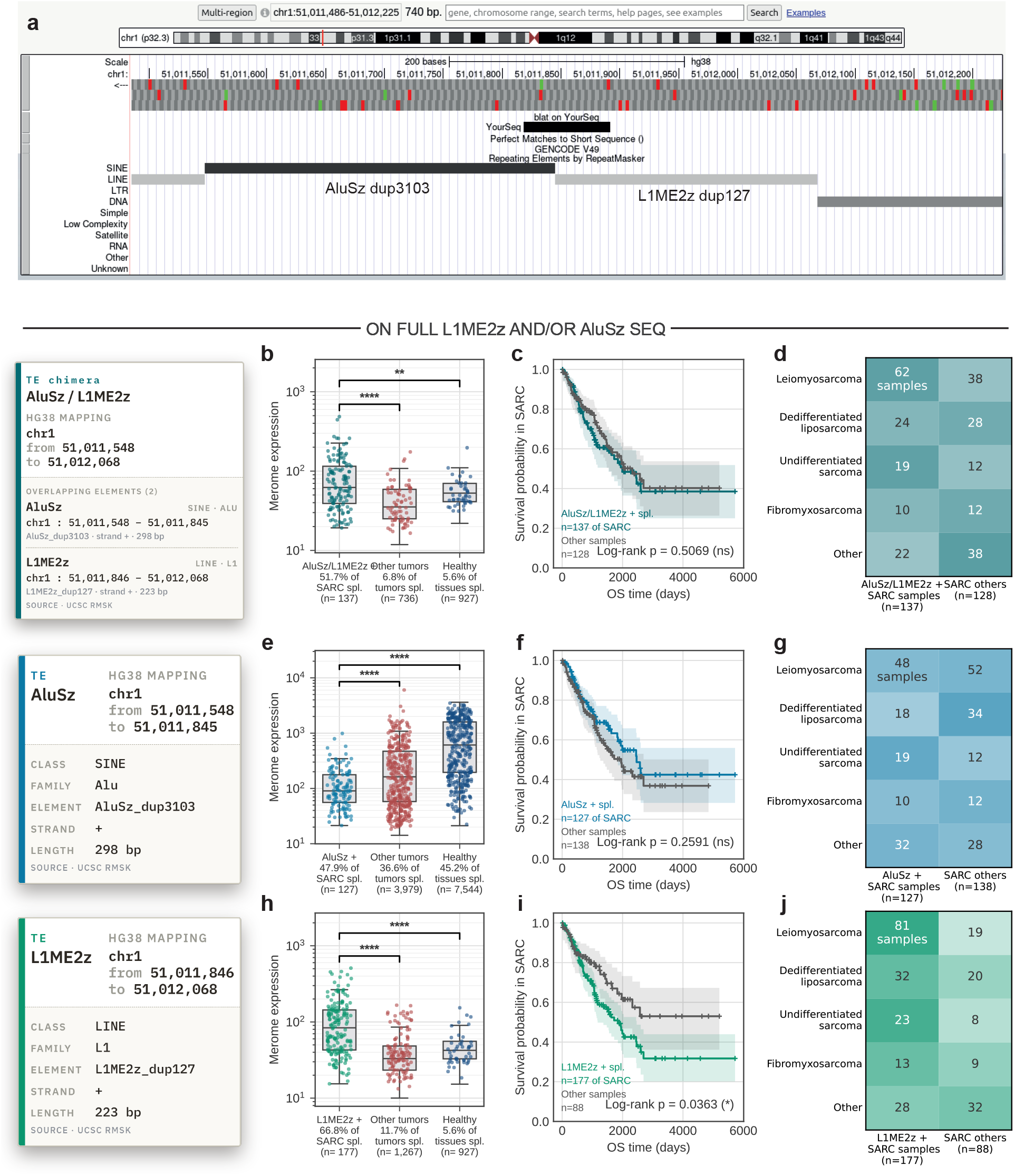
Only L1ME2z reproduces the read’s tumor-specificity and survival association; the adjacent AluSz does not. **a**, The read region (UCSC Genome Browser, chr1:51,011,498–51,012,225, hg38) overlaps two adjacent RepeatMasker elements, AluSz dup3103 (SINE/Alu) then L1ME2z dup127 (LINE/L1). Panels **b–j** test three full-length sequences: the AluSz/L1ME2z chimera (**b–d**; 521 bp), AluSz alone (**e–g**; 298 bp) and L1ME2z alone (**h–j**; 223 bp). **b, e, h,** Merome expression (sequence^+^ versus other tumors and healthy tissues; Mann–Whitney): the chimera (**b**; 51.7% of SARCs) and L1ME2z alone (**h**; 66.8% versus 5.6% of healthy tissues) are sarcoma-specific, whereas AluSz alone (**e**) is expressed across all samples, more in healthy tissues (45.2%) than other tumors (36.6%). **c, f, i,** Overall survival in SARC, sequence^+^ versus others: significant for L1ME2z alone (**i**; log-rank *p* = 0.0363), not for the chimera (**c**; *p* = 0.51) or AluSz alone (**f** ; *p* = 0.26). **d, g, j,** Sequence^+^ status by sarcoma histological subtype.

